# Structure-Functional-Selectivity Relationship Studies on A-86929 Analogs and Small Aryl Fragments toward Discovery of Biased D1 Agonists

**DOI:** 10.1101/2021.10.01.462758

**Authors:** Haoxi Li, Rosa Mirabel, Joseph Zimmerman, Ion Ghiviriga, Darian K. Phidd, Nicole Horenstein, Nikhil M. Urs

**Author notes:** Authors contributed equally (Haoxi Li, Rosa Mirabel).

## Abstract

Dopamine regulates normal functions such as movement, reinforcement learning, and cognition, and its dysfunction has been implicated in multiple psychiatric and neurological disorders. Dopamine acts through the D1- (D1R and D5R) and D2-class (D2R, D3R and D4R) of seven transmembrane receptors, and activates both G-protein- and β-arrestin-dependent signaling pathways, to mediate its physiological effects. Current dopamine receptor-based therapies are used to ameliorate motor deficits in Parkinson’s disease, or as antipsychotic medications for schizophrenia. These drugs show efficacy for ameliorating only some symptoms caused by dopamine dysfunction and are plagued by debilitating side-effects. Studies in primates and rodents have shown that shifting the balance of dopamine receptor signaling towards the arrestin pathway can be beneficial for inducing normal movement, while reducing motor side-effects such as dyskinesias, and can be efficacious at enhancing cognitive function compared to balanced agonists. Several structure-activity-relationship (SAR) studies have embarked on discovering β-arrestin-biased dopamine agonists, focused on D2 partial agonists, non-catechol D1 agonists, and mixed D1/D2R dopamine receptor agonists. Here, we describe an SAR study to identify novel D1R β-arrestin biased ligands using A-86929, a high-affinity D1R catechol agonist, as a core scaffold. Previously described and novel analogs of A-86929 were synthesized and screened *in vitro* for structure-functional-selectivity relationships (SFSR) studies to identify chemical motifs responsible for β-arrestin biased activity at both D1 and D2Rs. Most of the A-86929 analogs screened were G protein biased but none of them were exclusively arrestin-biased. Additionally, various catechol aryl fragments were designed and synthesized. Other compounds surveyed included hydroxyl and chloro analogs of dopamine to test for hydrogen bonding and ionic interactions. Some of these small molecular probes displayed weak bias towards the β-arrestin pathway. Continued in-depth SFSR studies informed by structure determination, molecular modeling, and mutagenesis studies will facilitate discovery of potent and efficacious arrestin-biased dopamine receptor ligands.

## BACKGROUND and significance

Dopamine signaling is central to the regulation of motor and cognitive function (Shiflett and Balleine, 2011; Smith and Graybiel, 2016; Wickens et al., 2007; Yin et al., 2006), and dopamine dysfunction is implicated in many disorders including Parkinson’s disease (PD) and in several psychiatric disorders (Jellinger and Korczyn, 2018; Shepherd, 2013; Surmeier et al., 2017; Wood and Ahmari, 2015). Dopamine mediates its physiological effects through two main G protein-coupled receptors (GPCRs), D1-class (D1R and D5R) and D2-class (D2R, D3R, D4R) of seven transmembrane receptors, which have been therapeutic targets for developing most antiparkinsonian and antipsychotic drugs. Current drug therapies for these disorders use dopamine receptor agonists or partial agonists which are balanced, i.e., they activate/inhibit both G protein and β-arrestin-dependent signaling pathways. Dopamine D1 or D2 agonist therapies for PD ameliorate motor symptoms but cause side-effects termed as dyskinesias, and do not improve cognitive function. For psychiatric disorders like schizophrenia, D2 receptor partial agonists/antagonists reverse the positive symptoms but do not correct the negative and cognitive symptoms, and have associated metabolic side-effects. We and others have previously shown that biasing dopamine receptor signaling towards the β-arrestin2 pathway can improve motor and cognitive function more efficaciously than balanced or G protein biased agonists (Allen et al., 2011; Park et al., 2015; Urs et al., 2015; Urs et al., 2016; Yang et al., 2018). Thus, targeting the β-arrestin pathway at dopamine receptors represents an excellent novel strategy to identify new pharmacotherapies for disorders like PD and schizophrenia.

Recently, biased signaling has gained a lot of traction for identifying novel, potent, efficacious, and pathway-selective compounds (Urban et al., 2007). Biased ligands have consistently been shown to selectively activate a single or subset of signaling cascade/s, in contrast to traditional balanced ligands that stimulate all possible responses upon binding to a receptor. We have recently shown that β-arrestin signaling is beneficial in PD models in promoting locomotion while reducing dyskinesias, through the action of D1 and D2Rs (Urs et al., 2015). Other groups have shown that weakly β-arrestin biased D1 agonists are much better than balanced D1 agonists in improving cognitive function (Yang et al., 2018). Previous SAR studies have identified D2 β-arrestin biased agonists that have a better therapeutic profile in mouse models of schizophrenia and also increase cortical interneuron firing significantly more than balanced agonists (Allen et al., 2011; Urs et al., 2016). However, the discovery of D1 β-arrestin biased agonists has remained challenging. One study identified potential G protein biased D1 agonists based on a benzazepine scaffold that are also antagonists for arrestin recruitment (Conroy et al., 2015). Recently, marginal β-arrestin biased D1 agonists based on the scaffold of dihydrexidine have been identified. These marginally arrestin biased agonists improve cortical coherence much better than G protein biased or balanced D1 agonists (Yang et al., 2018). More recent SAR studies have identified high affinity G protein biased D1 agonists based on non-catechol scaffolds (Gray et al., 2018; Martini et al., 2019a; Martini et al., 2019b; Wang et al., 2019), and these G protein biased agonists robustly enhanced locomotor activity in mice. We have previously embarked on studies to identify D1 or D2 arrestin biased agonists based on the scaffold of mixed dopamine agonist Apomorphine (Park et al., 2020). Our studies identified some marginally arrestin biased D1 or D1/D2R agonists. Thus far, identifying high affinity arrestin biased D1 agonists has remained challenging.

We therefore focused our efforts on parent drug scaffolds that are high affinity and efficacy D1 agonists that have been clinically tested for motor and cognitive function. We used the D1 agonist ABT-431, the diacetyl prodrug of A-86929, that has been tested in the clinic and has been shown to be efficacious in ameliorating PD motor symptoms (Michaelides et al., 1995; Rascol et al., 1999). However, ABT-431 also induced dyskinesias to the same extent as L-DOPA therapy (Rascol et al., 2001). A-86929 analogs have been synthesized and studied before but these analogs were not tested for their ability to activate G protein versus β-arrestin pathways. Our goal was to synthesize these previously published analogs and some novel analogs and test their ability to activate G protein and β-arrestin signaling at D1 and D2Rs. In addition to A-86929 analogs we tested several smaller fragments to mimic interactions within the binding pocket of dopamine receptors and have been hypothesized to be important for G protein or arrestin activity (Gray et al., 2018; Zhuang et al., 2021a).

## RESULTS

### 1. Target compound syntheses

To determine whether modifying various structural elements of A-86929 would result in biased compounds favoring either cAMP or β-arrestin2 signaling, we focused on substitutions on the thiophene or the nitrogen center. We also explored different configurations at the two stereogenic centers since we and others have shown differential activity between the enantiomers (Michaelides et al., 1997; Park et al., 2020).

#### 1.1. Synthesis of A-86929 analogs

The approaches used to synthesize analogs of A-86929, compounds **1a**-**1i**, and its two enantiomers are presented in Schemes 1 and 2. The route utilizes the reported approach (Hajra and Bar, 2011; Sodergren et al., 1997) featuring Evans aziridination and in-situ cyclization to the target tetracyclic compounds. The route differs in the approach to alkenes **6a-6c**, because we wanted to have a latent center on the thiophene for coupling reactions, which we considered might have been incompatible with the reported route utilizing organoboranes. In our hands the aziridination chemistry proceeded in low (unoptimized) yields, but the enantiomeric enrichment with the Box ligands was moderate, ranging from 80 to 91%. We found the Br substituent on the thiophene ring was an excellent partner for Pd mediated coupling to produce a number of probative compounds and their enantiomers. (Scheme 2).

**Scheme 1.**
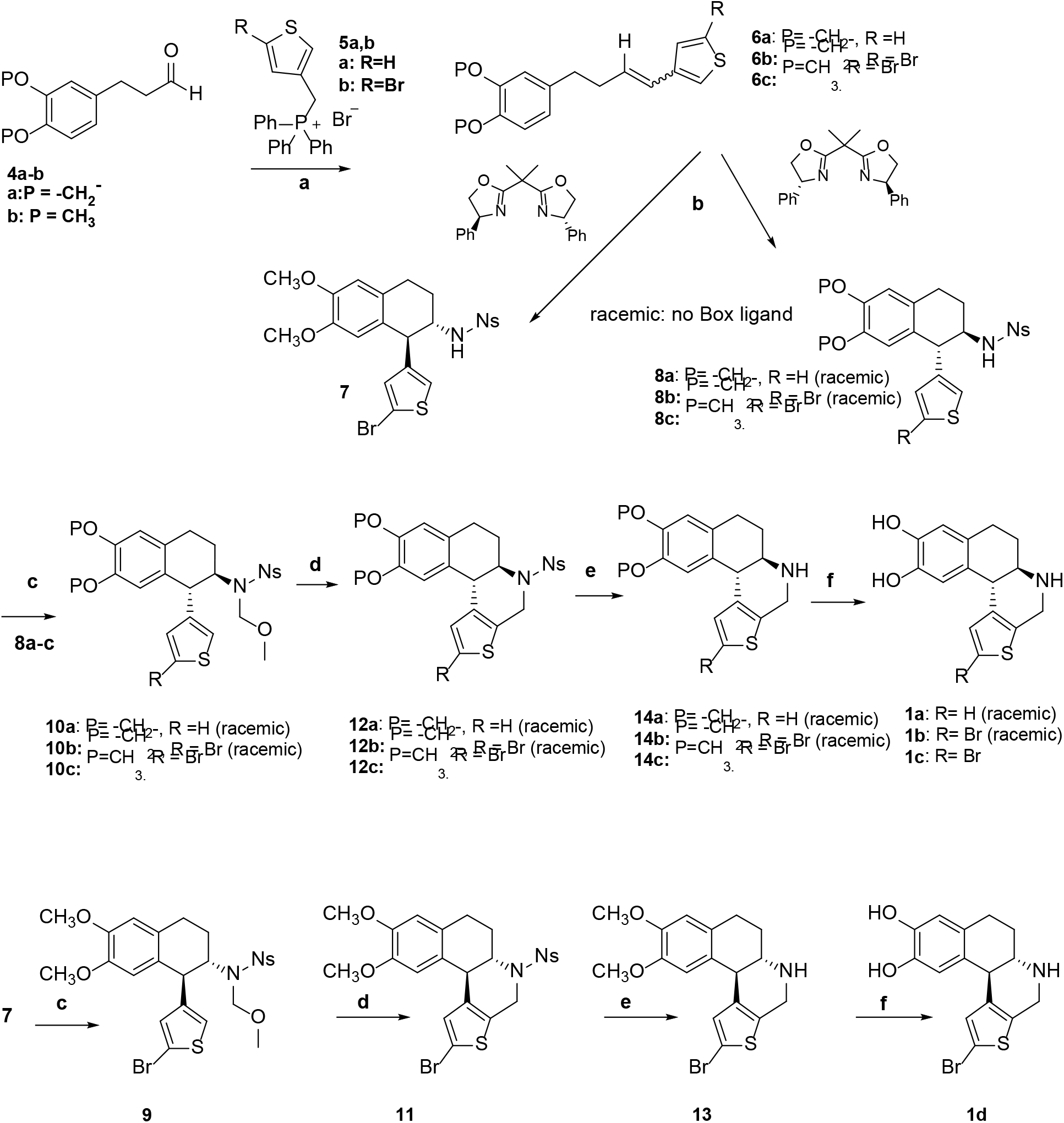
Synthesis of A-86929 analogs **1a-d**^a^ ^a^Reaction conditions: (a) DBU, DCM, (61-73 %); (b) Cu(OTf)_2_, PhI=NNs, DCM, 3 or 4 Å molecular sieves, DCM, (6-16 %); (c) NaH, MOMCl, THF, (75-78 %); (d) TMSOTf, DCM, (64-93 %); (e) PhSH, K_2_CO_3_, CH_3_CN, DMSO (85 %-quant); (f) BCl_3_ or BBr_3_, DCM, (67 %-quant).

**Scheme 2.**
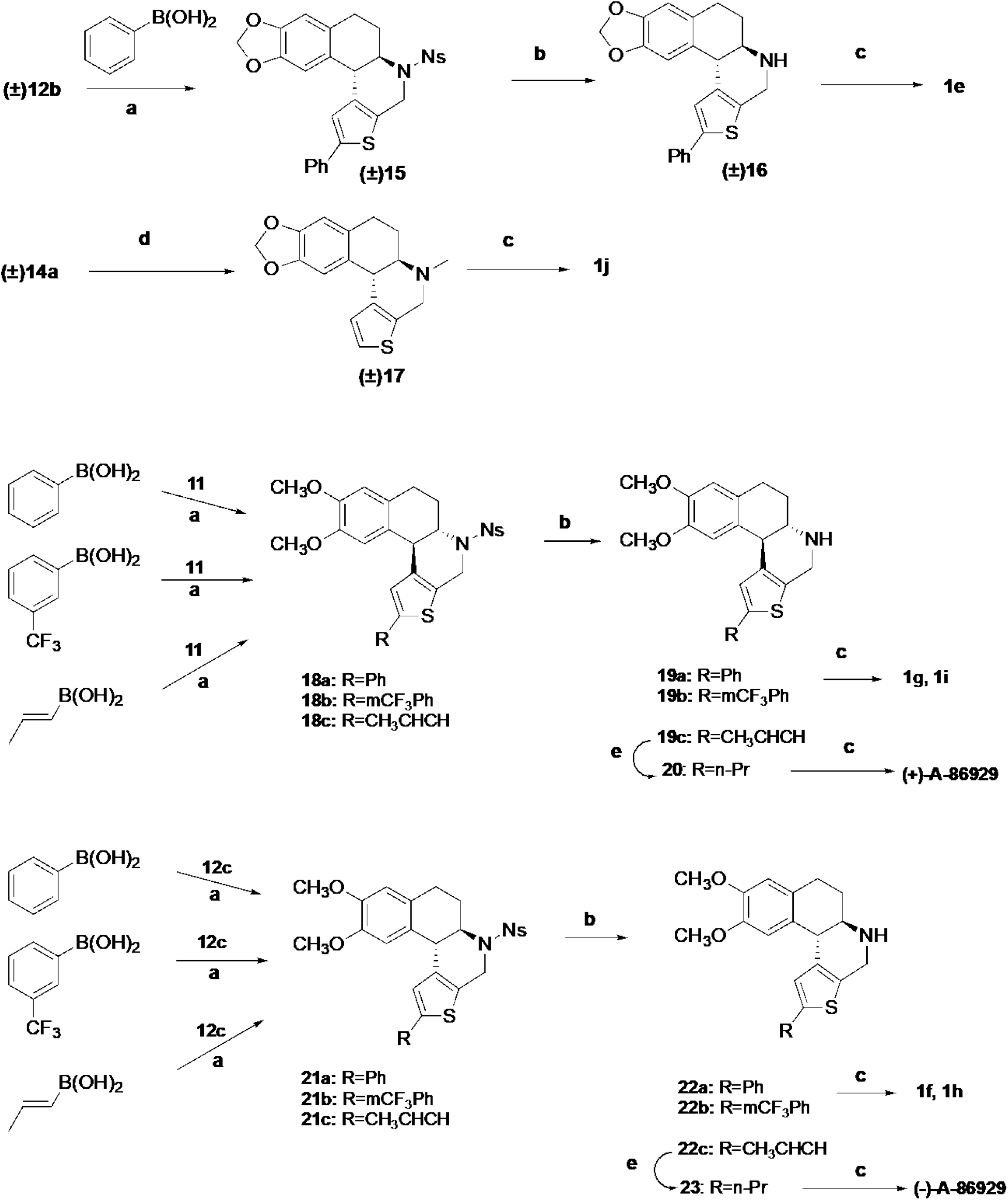
Synthesis of A-86929 analogs **1e-1j** and both antipodes of A-86929.^a^ ^a^Reaction conditions: (a) Pd(PPh_3_)_4_, Na_2_CO_3_, DME/H_2_O, (53-91 %); (b) PhSH, K_2_CO_3_, CH_3_CN, DMSO (85 %-quant); (c) BCl_3_ or BBr_3_, DCM, (26-69%); (d) HCHO, AcOH, NaBH_3_CN, CH_3_CN, (39 %); (e) H_2_, Pd-C, MeOH, (60-96%).

We also synthesized some acyclic analogs of A-86929, compounds **1l** and **1m** in racemic form via compound **24**, (Michaelides et al., 1997) in good yields. (Scheme 3)

**Scheme 3.**
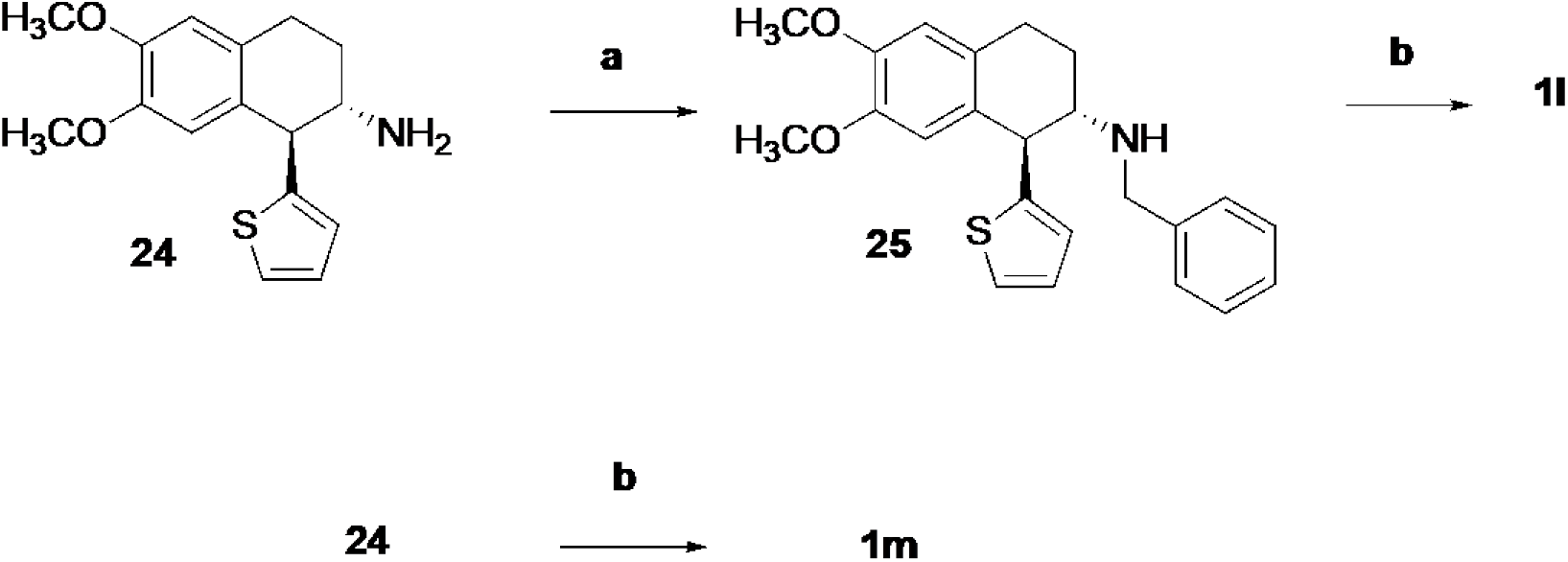
Synthesis of acyclic A-86929 analogs **1l, 1m**^a^ ^a^Reaction conditions: (a) PhCHO, NaBH_3_CN, AcOH/MeOH, (86 %); (b) BBr_3_, DCM, (27-97 %)

#### 1.2. Evaluation of A-86929 analogs in cell-based assays

All compounds were evaluated for their activities for stimulating G-protein signaling using a cAMP level-dependent chemiluminiscent sensor (GloSensor assay, Promega) and β-arrestin2 recruitment (Bioluminiscence Resonance Energy Transfer or BRET assay) at D1 and D2 receptors using standard *in vitro* dose-response assays (see Methods section). The results in the tables show the potency (EC_50_, pEC_50_) and efficacy (E_max_) values calculated using the sigmoidal dose-response function in GraphPad Prism 9.0. The percent response values were normalized to dopamine. As shown in Table 1, ABT-431, which is the prodrug (acetylated catechol groups) of A-86929, is a high affinity D1 agonist with nanomolar potency at the Gαs pathway, consistent with previous observations (Michaelides et al., 1997). However, ABT-431 has low micromolar potency at the arrestin pathway. As expected, at D2R, ABT-431 has low micromolar potency at both pathways. A-86929 which is the non-acetylated form of ABT-431 is more potent than ABT-431 at D1Rs in vitro, with the (-)-enantiomer more potent at both pathways compared to the (+)-enantiomer. At D2Rs, both enantiomers of A-86929 did not affect the G protein pathway but the (-)-enantiomer enhanced potency and efficacy at the β-arrestin pathway, compared to ABT-431. Variation of substitution at the R1 position of A-86929 with either H (unsubstituted) **1a** or bromine **1b** enhanced potency and efficacy at both pathways at D1Rs and D2Rs compared to ABT-431 (Figure 2). Compared to A-86929, these substitutions, enhanced potency and efficacy at the D1 but only potency (>10-fold) at the D2 G protein pathway. In contrast, these substitutions enhanced potency and efficacy compared only to the (-) enantiomer of A86929 at both D1/D2R arrestin pathways, but showed increased or decreased efficacy with marginal changes to potency, compared to the (+) enantiomer at D1 or D2R arrestin pathways, respectively (Table 1).

**Table 1.** Agonist activities of A-86929 analogs EC_50_ and E_max_ values are from three independent experiments performed in duplicate or triplicate. E_max_ values are calculated as % response normalized to dopamine. N.D not determined.

| Cmpd | β-arrestin | | | cAMP | | |
| --- | --- | --- | --- | --- | --- | --- |
| | EC<sub>50</sub> (nM) | pEC<sub>50</sub> ± SEM | E<sub>max</sub> (%) ± SEM | EC<sub>50</sub> (nM) | pEC<sub>50</sub> ± SEM | E<sub>max</sub> (%) ± SEM |
| **D1R** |  |  |  |  |  |  |
| DA | 319.70 | 6.5±0.11 | 90.18±3.73 | 1.20 | 8.9±0.1 | 81.51±3.12 |
| ABT-431 | 3200.00 | 5.49±0.07 | 88.17±3.06 | 1.05 | 8.98±0.08 | 108.9±3.64 |
| (+)-A-86929 | 2800.00 | 5.56±0.15 | 94.12±6.55 | 0.26 | 9.6±0.87 | 91.33±9.2 |
| (-)-A-86929 | 607.00 | 6.22±0.16 | 95.46±6.88 | 0.11 | 9.96±1.13 | 93.91±7.1 |
| 1a | 797.70 | 6.10±0.15 | 113.31±7.66 | 0.04 | 10.4±0.11 | 115.33±5.32 |
| 1b | 515.67 | 6.29±0.14 | 137.89±8.41 | 0.05 | 10.3±0.19 | 111.34±8.51 |
| 1c | 1092.20 | 5.96±0.15 | 136.51±9.35 | 0.45 | 9.35±0.56 | 72.73±10.9 |
| 1d | 2632.50 | 5.58±0.18 | 52.83±9.7 | 0.69 | 9.16±0.51 | 52.83±9.7 |
| 1e | >10000 | N.D | N.D | 2.70 | 8.6±0.21 | 74.12±6.1 |
| 1f | >10000 | N.D | N.D | 3.55 | 8.45±0.41 | 50.43±8.4 |
| 1g | >10000 | N.D | N.D | 4.73 | 8.32±0.7 | 71.5±12.5 |
| 1h | >10000 | N.D | N.D | 3.77 | 8.42±0.49 | 67.7±13.43 |
| 1i | >10000 | N.D | N.D | 63.80 | 7.2±0.21 | 77.0±6.7 |
| 1j | >10000 | N.D | N.D | 1.20 | 8.9±0.23 | 63.04±6.24 |
| 1k | >10000 | N.D | N.D | 114.30 | 6.94±0.41 | 45.9±8.03 |
| 1l | >10000 | N.D | N.D | 11.30 | 7.95±0.55 | 36.5±8.5 |
| 1m | 251.2 | 6.6±1.1 | 3.7±1.67 | 36 | 7.44±0.3 | 58.4±6.8 |
| **D2R** |  |  |  |  |  |  |
| DA | 54.10 | 7.3±0.09 | 90.29±3.23 | 0.71 | 9.2±0.22 | 66.23±6.78 |
| ABT-431 | 3000.00 | 5.53±0.2 | 22.99±2.15 | 2986.60 | 5.52±0.11 | 105.33±5.41 |
| (+)-A-86929 | 2074.00 | 5.68±0.25 | 38±4.45 | 2937.30 | 5.53±0.21 | 140.74±13.69 |
| (-)-A-86929 | 270.20 | 6.57±0.12 | 79.7±3.83 | 7474.6 | 5.13±0.26 | 122.78±17.82 |
| 1a | 169.93 | 6.77±0.05 | 55.17±1.17 | 115.30 | 6.94±0.42 | 56.92±10.19 |
| 1b | 581.20 | 6.24±0.09 | 60.34±2.27 | 197.50 | 6.70±0.19 | 73.49±8.02 |
| 1c | 279.60 | 6.55±0.12 | 70.3±3.41 | 957.16 | 6.02±0.22 | 78±8.14 |
| 1d | 5624.60 | 5.25±0.15 | 50.63±4.02 | 7360.20 | 5.13±0.32 | 123±21.8 |
| 1e | >10000 | N.D | N.D | 60.60 | 7.2±0.3 | 70±8.8 |
| 1f | 5819.40 | 5.24±0.1 | 85.51±4.64 | 1097.30 | 5.96±0.25 | 96.3±11.2 |
| 1g | >10000 | N.D | N.D | 1373.50 | 5.86±0.21 | 97.9±9.6 |
| 1h | 4840.30 | 5.32±0.13 | 73.05±4.69 | 1654.20 | 5.78±0.35 | 70.92±11.7 |
| 1i | >10000 | N.D | N.D | 8378.00 | 5.08±0.26 | 93.1±14.15 |
| 1j | 7484 | 5.13±0.43 | 73.4±17.8 | 154.60 | 6.8±0.28 | 76.7±9.6 |
| 1k | >10000 | N.D | N.D | 600.4 | 6.22±1.1 | 36.2±16.9 |
| 1l | 8174 | 5.77±0.6 | 8.62±3.15 | >10000 | N.D | N.D |
| 1m | 5440 | 5.3±0.12 | 27±1.8 | 847.2 | 6.1±0.5 | 76.44±16.4 |

**Figure 1.**
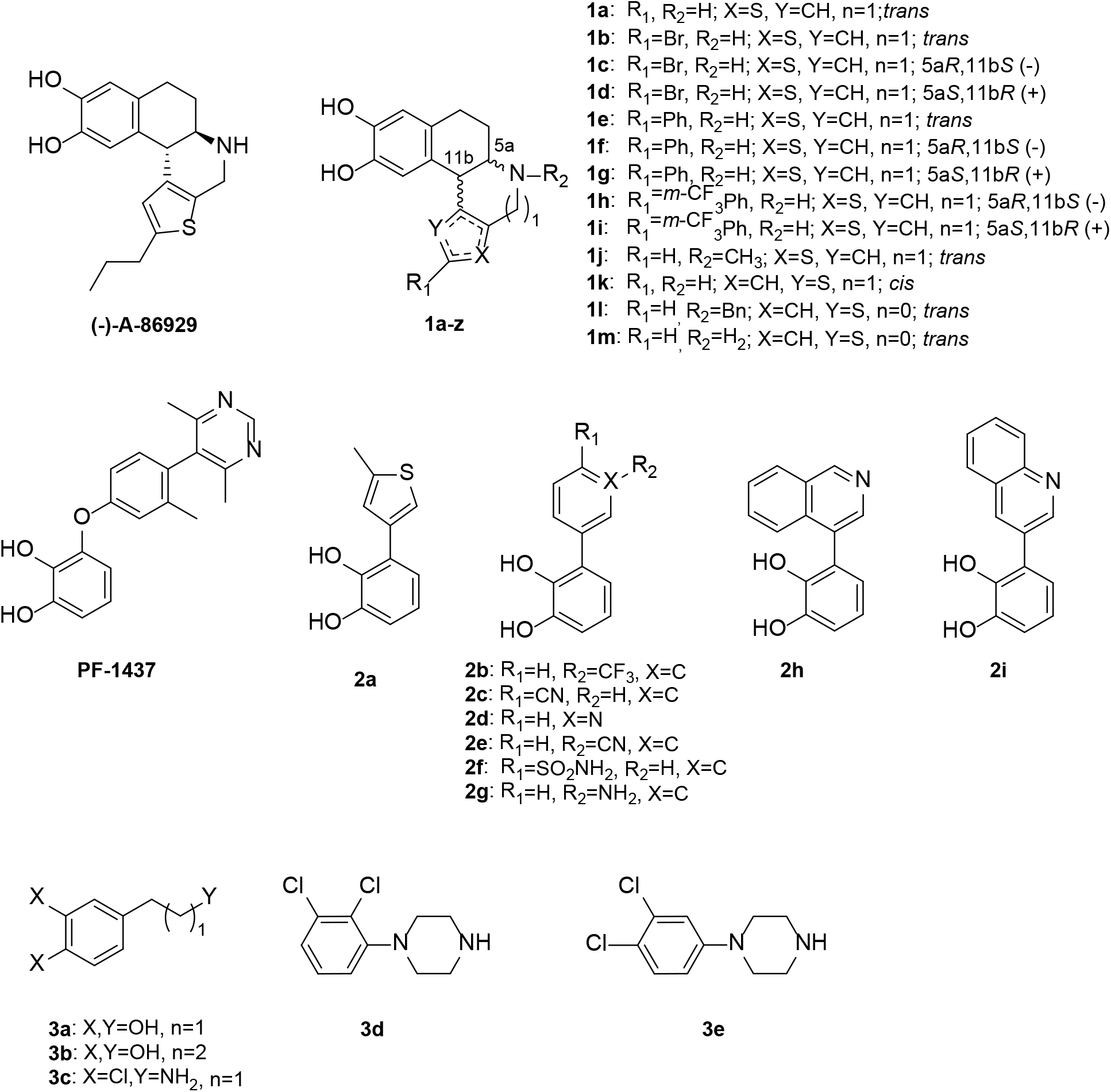
Structures of compounds we used in these studies. Compound syntheses are summarized in Schemes.

**Figure 2.**
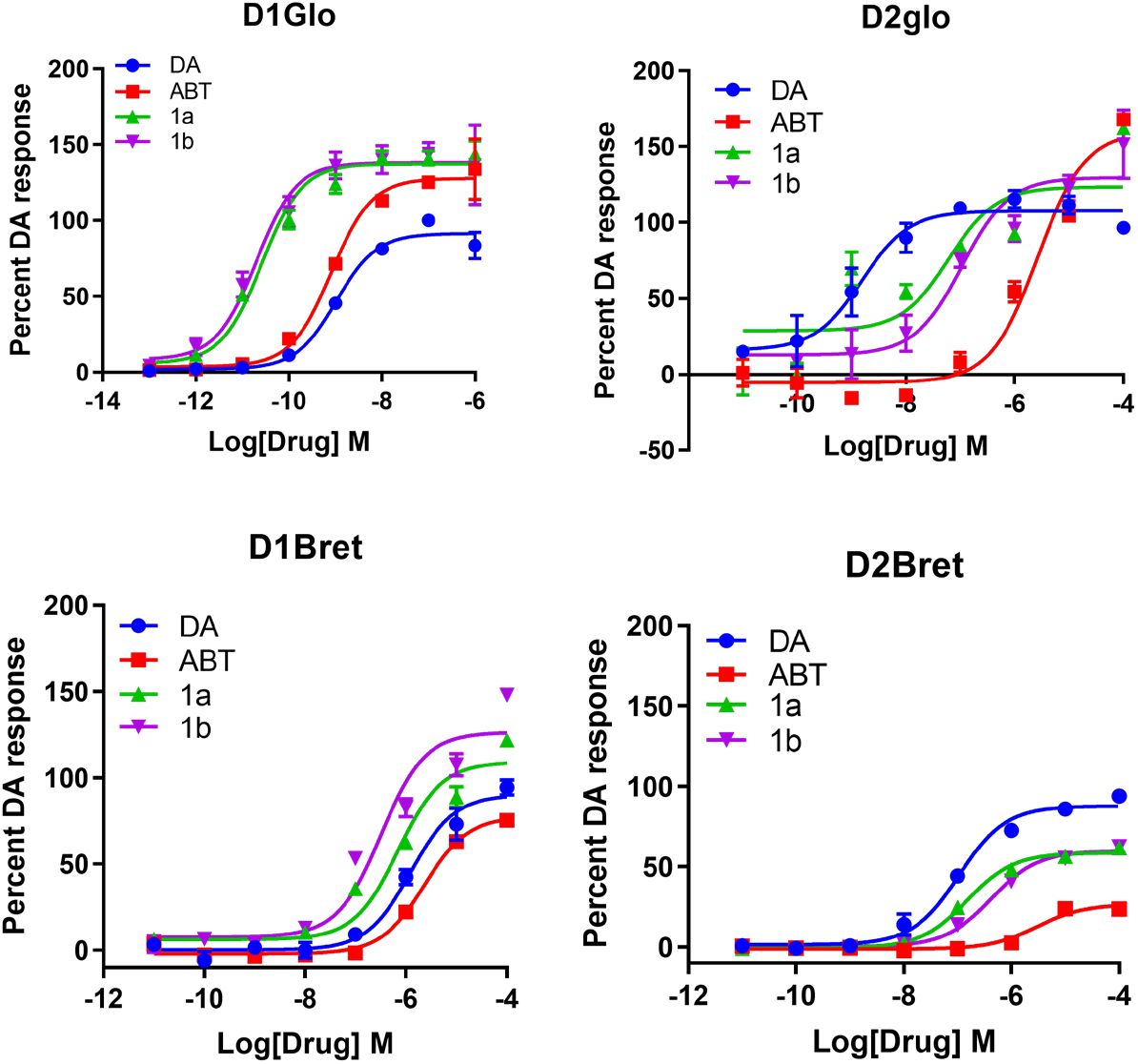
Dose response curves comparing ABT-431 and substituted analogs. *In vitro* GloSensor and BRET agonist assays at D1Rs and D2Rs for substituted analogs (**1a, 1b**) in HEK-293 cells. Data are presented as percent of the total DA response (mean ± SEM).

Addition of a bulky phenyl or phenyl-CF_3_ group at this position completely abolished β-arrestin activity at D1R, and reduced G protein activity at both D1 and D2Rs. We further calculated signaling bias of these compounds for the G protein vs β-arrestin pathways at D1 and D2Rs using the operational model. We calculated the ΔΔLog (Tau/K_A_) values of the compounds in Table 1, and as shown in Figure 3, A-86929 and its analogs are G protein biased at D1Rs. However, at D2Rs, A-86929 (+) and (-) to a larger extent and its analog 1c shows arrestin bias, but most of the remaining analogs are G protein biased.

**Figure 3.**
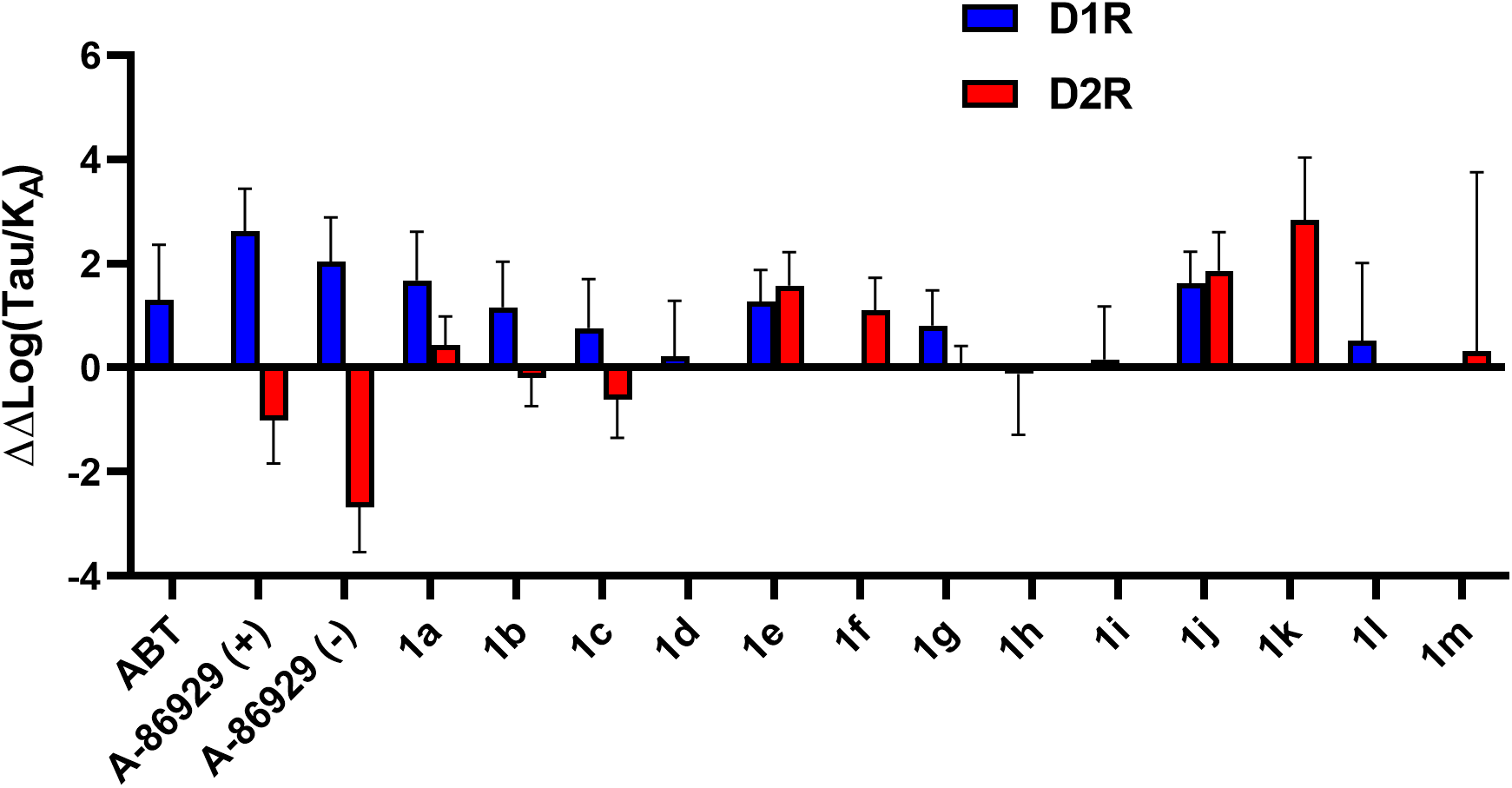
Bias calculations for compounds in Table 1. ΔΔLog(Tau/K_A_) values of compounds were calculated for G protein and β-arrestin pathways at D1 and D2Rs. Negative and positive values represent arrestin and G protein bias, respectively.

### 2. Synthesis and signaling pathway evaluation of small fragments

Since most of the modifications to the A-86929 scaffold did not yield any potent and efficacious D1R β-arrestin biased compounds, we altered our SFSR approach. Unlike dopaminergic agonists, non-catechol scaffolds mainly rely on contacts with ECL2 (e.g., L190^ECL2^) to activate G protein signaling pathway on D1R, rather than serine residues on TM5 (e.g., S198^5.42^, S202^5.46^) and D103^3.32^ on TM3 (Gray et al., 2018) (Ballesteros-Weinstein numbering rules were applied to the superscripted GPCRs numbering (Ballesteros and Weinstein, 1995)). Catechol modification on a G protein biased non-catechol ligand (PF-8871) yielded a hybrid compound (PF-1437) with a balanced signaling profile. This indicates that interactions between the catechol moiety and serine residues on TM5 might be advantageous for β-arrestin signaling activation. Recent cryo-electron microscopy (cryo-EM) studies with D1 and D2Rs have shown that the specificity of drugs for D1 vs D2 binding pockets arise from the interactions of these compounds with the ECL2 (Xiao et al., 2021; Zhuang et al., 2021a). Therefore, our second goal was to develop small fragments based on this hybrid compound by maintaining the catechol motif while truncating the size of the molecule. This strategy potentially maintains interactions between the catechol and TM5 serine residues for β-arrestin activation, and simultaneously reduces contacts with ECL2 to decrease G protein signaling. We hypothesized that such smaller fragments will give us insights into potential interactions that are important for G protein vs β-arrestin activity.

We synthesized the 2-aryl substituted catechols **2a-2i** as presented in Scheme 4, with fair to excellent yields utilizing Pd-mediated coupling reactions.

**Scheme 4.**
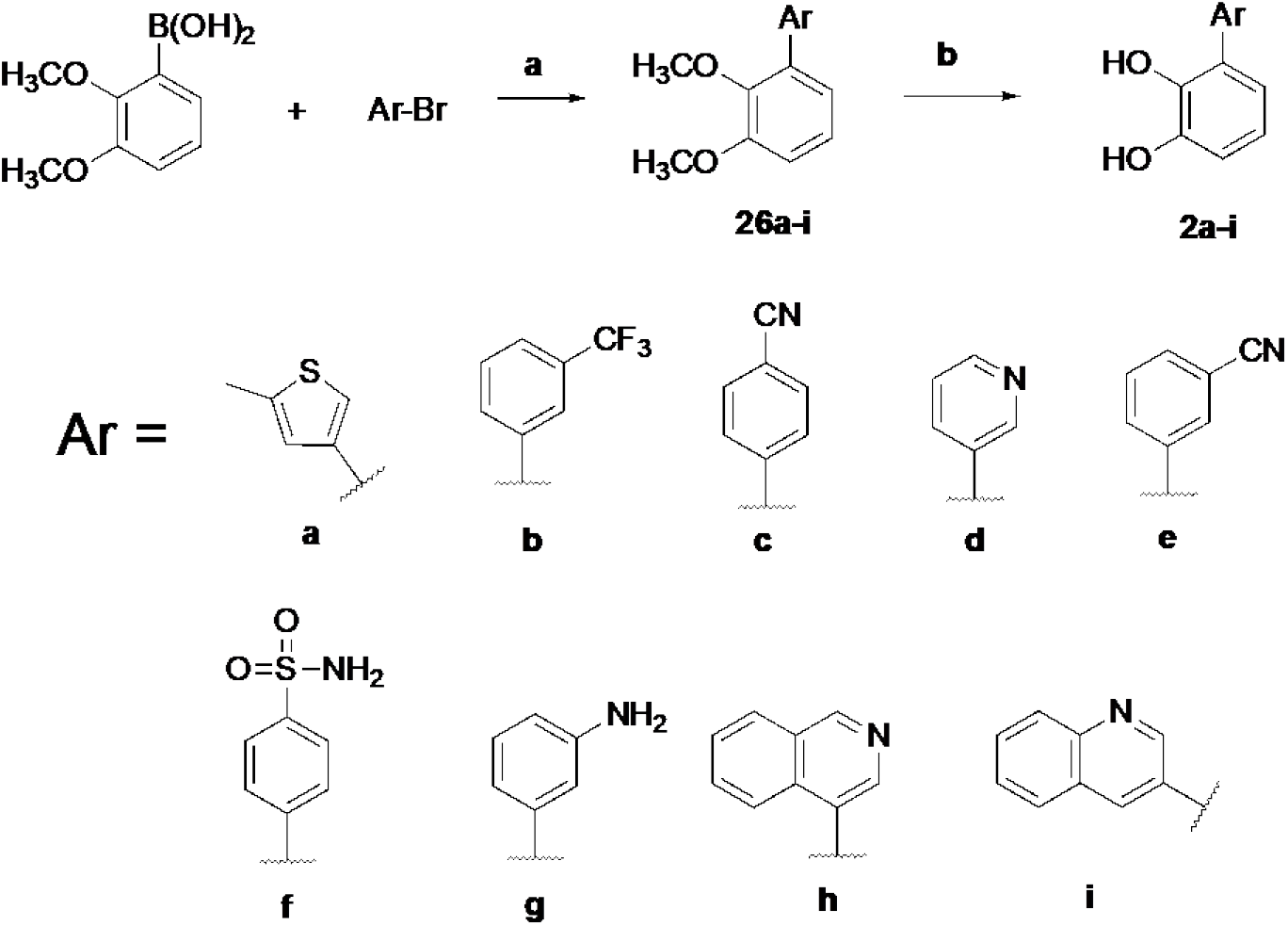
Synthesis of 2-aryl catechols.^a^ ^a^Reaction conditions: (a) Pd(PPh_3_)_4_, Na_2_CO_3_, DME/H_2_O, (36-80 %); (b) BBr_3_, DCM, (27-97 %).

As seen in Table 2, these modifications significantly reduced activity at both pathways at D1 and D2Rs. However, **2d, f, g** and **h** showed certain degree of bias for the β-arrestin pathway only at D1Rs, even though these compounds have very low efficacy. Conversely, **2a** and **2c** showed certain degree of bias for the arrestin pathway at D2Rs but not D1Rs (Figure 4). In addition to the 2-aryl fragments we also tested some 3-aryl fragments commercially purchased from Enamine. Although these fragments generally showed weak to no activity at D1/D2Rs, some fragments showed interesting properties. The fragment with the meta methoxy group (Z4642446630) was marginally D1R arrestin biased, but if the methoxy was para (EN30010864588) or ortho (Z4642443452), these analogs were D1R G protein biased. However, these fragments were weak partial agonists (Table S1).

**Table 2.** Agonist activities of catechol fragments EC_50_ and E_max_ values are from three independent experiments performed in duplicate or triplicate. E_max_ values are calculated as % response normalized to dopamine. ND - not detectable, N.D. not determined.

| Cmpd | β-arrestin | | | cAMP | | |
| --- | --- | --- | --- | --- | --- | --- |
| | EC<sub>50</sub> (nM) | pEC<sub>50</sub> ± SEM | E<sub>max</sub> (%) ± SEM | EC<sub>50</sub> (nM) | pEC<sub>50</sub> ± SEM | E<sub>max</sub> (%) ± SEM |
| **D1R** |  |  |  |  |  |  |
| DA | 319.70 | 6.5±0.11 | 90.18±3.73 | 1.20 | 8.9±0.1 | 81.51±3.12 |
| 2a | 6895 | 5.16±0.32 | 30.31±5.6 | 130 | 6.88±0.59 | 19.7±5.54 |
| 2b | 5.2 | 8.28±1.21 | 6.13±3.11 | 98 | 7.01±0.77 | 16.62±6.06 |
| 2c | >10000 | N.D | N.D | 0.15 | 9.81±1.12 | 16.30±9.94 |
| 2d | 53 | 7.28±1.7 | 2.6±1.8 | >10000 | N.D | N.D |
| 2e | >10000 | N.D | N.D | 3605.2 | N.D | N.D |
| 2f | 51.4 | 7.3±0.8 | 4.6±1.5 | >10000 | N.D | N.D |
| 2g | 7.33 | 8.13±0.9 | 4±1.6 | >10000 | N.D | N.D |
| 2h | 689.1 | 6.2±0.6 | 8.1±2.1 | >10000 | N.D | N.D |
| 2i | >10000 | N.D | N.D | 590.2 | 6.23±0.9 | 9.7±3.8 |
| **D2R** |  |  |  |  |  |  |
| DA | 54.10 | 7.3±0.09 | 90.29±3.23 | 0.71 | 9.2±0.22 | 66.23±6.78 |
| 2a | 1681.7 | 5.09±0.64 | 13.9±3.68 | >10000 | N.D | N.D |
| 2b | >10000 | N.D | N.D | >10000 | N.D | N.D |
| 2c | 31.69 | 7.5±2.02 | 1.61±1.27 | >10000 | N.D | N.D |
| 2d | ND | ND | ND | >10000 | N.D | N.D |
| 2e | ND | ND | ND | >10000 | N.D | N.D |
| 2f | ND | ND | ND | >10000 | N.D | N.D |
| 2g | ND | ND | ND | >10000 | N.D | N.D |
| 2h | ND | ND | ND | >10000 | N.D | N.D |
| 2i | ND | ND | ND | >10000 | N.D | N.D |

**Figure 4.**
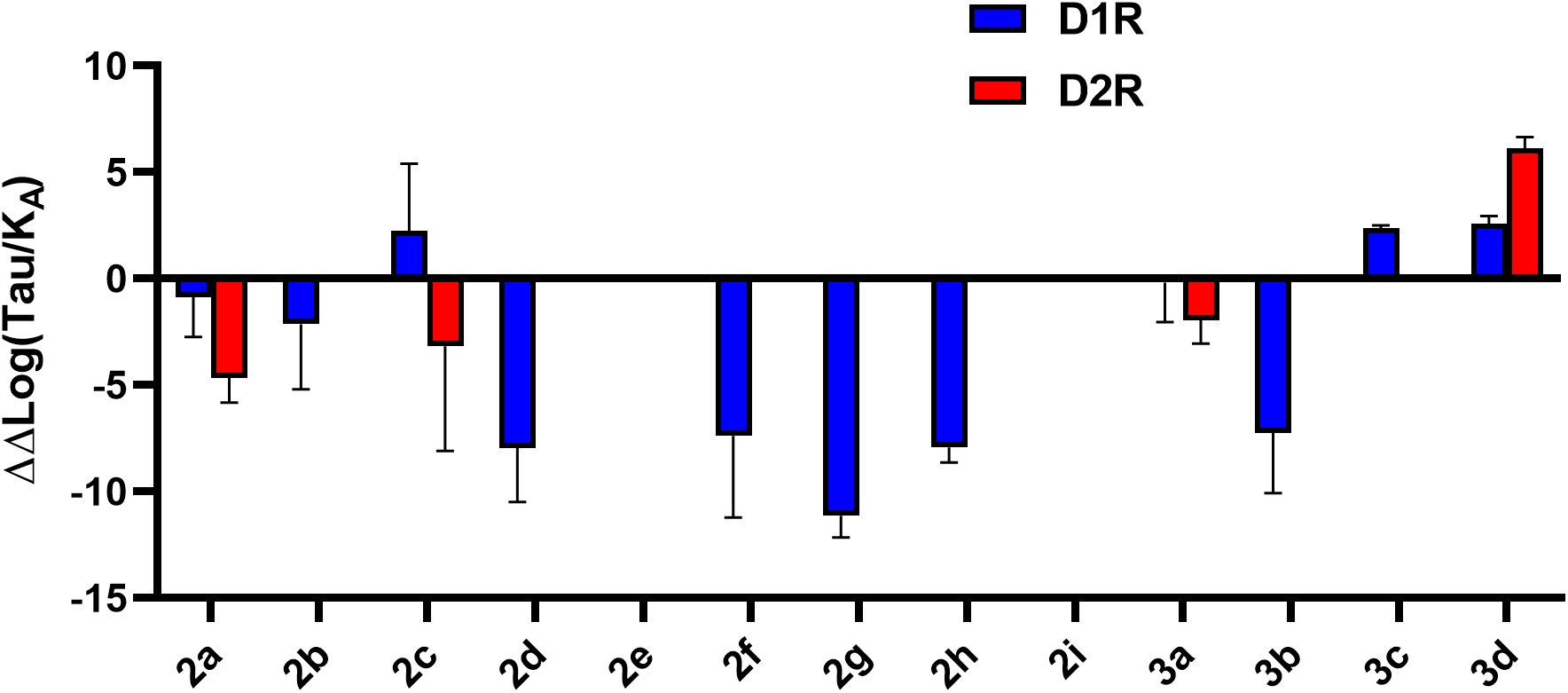
Bias calculations for compounds in table 2. ΔΔLog (Tau/KA) values of compounds were calculated for G protein and β-arrestin pathways at D1 and D2Rs. Negative and positive values represent arrestin and G protein bias, respectively.

Finally, we tested a small group of low molecular weight fragment compounds in which we used dopamine analogs that were probative for substitution of the amino group charge for H-bonding (Figure 1, compounds **3a**,**b**) or replacement of the catechol oxygen with chlorine, (Figure 1, compounds **3c**, **3d**,**e**). Their activities are shown in Table 3. Most of these modifications had a general inhibitory effect at all pathways, with a tendency to be G protein biased (Figure 4), with only fragments with hydroxyl groups (**3a**, **3b**) showing some degree of bias for D2R or D1R arrestin, respectively. Compound 3d was a full agonist for the G-protein pathway at D2R, and the rest of the fragments were partial agonists.

**Table 3.** Agonist activities of dopamine-like probes EC_50_ and E_max_ values are from three independent experiments performed in duplicate or triplicate. E_max_ values are calculated as % response normalized to dopamine. ND - not detectable, N.D. not determined.

| Cmpd | β-arrestin | | | cAMP | | |
| --- | --- | --- | --- | --- | --- | --- |
| | EC<sub>50</sub> (nM) | pEC<sub>50</sub> ± SEM | E<sub>max</sub> (%) ± SEM | EC<sub>50</sub> (nM) | pEC<sub>50</sub> ± SEM | E<sub>max</sub> (%) ± SEM |
| D1R |  |  |  |  |  |  |
| 3a | 228.9 | 6.64±0.65 | 11.46±3.05 | 1090 | 5.96± 0.46 | 60.57± 16.66 |
| 3b | 534.2 | 6.3±1.2 | 3.5±1.9 | 5171 | 5.3±0.8 | 21±8.13 |
| 3c | ND | ND | ND | 4584 | 5.34±0.1 | 24.43±1.2 |
| 3d | ND | ND | ND | 580 | 6.24±0.15 | 5.03±0.32 |
| 3e | ND | ND | ND | ND | ND | ND |
| D2R |  |  |  |  |  |  |
| 3a | 5260 | 5.28±0.57 | 9.98±2.96 | >10000 | N.C | N.C |
| 3b | >10000 | N.D | N.D | >10000 | N.D | N.D |
| 3c | ND | ND | ND | >10000 | N.C | N.C |
| 3d | ND | ND | ND | 698.1 | 6.12±0.13 | 138.42±8.03 |
| 3e | ND | ND | ND | >10000 | N.D | N.D |

## DISCUSSION

In this current SFSR study, we used the dopamine D1 agonist A-86929 and PF-1437 as our parent scaffolds to identify novel arrestin biased D1R analogs. L-DOPA and mixed dopamine agonist therapies are efficacious in reversing PD motor symptoms but have associated side-effects such as dyskinesias. ABT-431, the diacetyl prodrug of A-86929, was synthesized as the first D1 agonist for PD and showed great clinical efficacy in PD but also induced dyskinesias, and did not undergo any further SAR or clinical studies. The original A-86929 analogs were solely tested for cAMP effects at D1 and D2Rs since the β-arrestin pathway was not yet identified as a potential signaling effector for dopamine receptors. Since the discovery of arrestin-dependent effects at dopamine receptors and evidence showing a beneficial effect of this pathway for motor symptoms of PD, SAR studies have sought to discover such arrestin-biased dopamine receptor agonists. Our goal in this study was to screen previously known and novel analogs of A-86929 for their ability to selectively activate the β-arrestin pathway at D1R and potentially at D2R. Our approach was to systematically make substitutions on different regions of the A-86929 scaffold and perform cell-based screening assays for G protein versus β-arrestin recruitment activity at D1 and D2Rs.

We first generated previously synthesized analogs of A-86929 (Michaelides et al., 1995; Michaelides et al., 1997) and tested their *in vitro* activity. Consistent with the previous studies, ABT-431 has low nanomolar potency at D1Rs but high micromolar potency at D2Rs. Since A-86929 was originally found to have weak activity at D2R using a G protein-based assay, we hypothesized that this compound could be arrestin biased at D2Rs, since activity at both receptors is needed under hypodopaminergic conditions. As expected, A-86929 did show arrestin bias at D2Rs, although it was a weak partial agonist. Replacing the propyl group (R1 position) on the thiophene of A-86929 with smaller groups such as H, increased potency at both D1 (picomolar) and D2Rs (low micromolar) at the G protein pathway. This is consistent with recent cryo-EM structures of D1 and D2Rs showing that pure D1 agonists protrude towards the ECL2 making them selective for D1Rs (Xiao et al., 2021; Zhuang et al., 2021a).

To explore binding conformations of these molecules in D1 and D2Rs, we implemented molecular docking with recent cryo-EM structures of D1R (Xiao et al., 2021; Zhuang et al., 2021a; Zhuang et al., 2021b). The binding poses of A-86929 and its analogs in D1R show common interactions of dopaminergic ligands and D1R, for example, hydrogen bonding interactions between the catechol motif and S198^5.42^, S202^5.46^ or N292^6.55^ residues, ionic salt bridge of the protonated amine and D103^3.32^, and π-π interactions with F288^6.51^ or F289^6.52^ (Fig. 5a-b). A previous study has shown that enantiomeric isomers of A-86929 analogs have different potency and efficacy at D1Rs (Michaelides et al., 1997), which is consistent with our results. Similarly, it has been reported that *R* and *S* enantiomers of benzazepine type agonists and antagonists (e.g., SKF-83959 and SCH-23390) also displayed different affinities towards D1R, with the *R* enantiomer as the primary contributor (Felsing et al., 2019; Zhang et al., 2009). Based on the docking results, we found that the 5a*R*, 11b*S* (-) enantiomers of A-86929 analogs can place the substituents on the thiophene into an extended binding pocket (EBP), surrounded by S188^ECL2^, S189^ECL2^, L190^ECL2^, A195^5.39^, F288^6.51^, N292^6.55^, L295^6.58^, P296^6.59^ and F313^7.35^ on the extracellular site of TM5, TM6, TM7 and ECL2. However, there is not as much space on the extracellular site of TM3 and ECL2 to accommodate substituents on the other enantiomer (Fig. 6a-d), which engenders steric hindrance, especially for bulky substitutions. This might explain such differences in potency of the enantiomers. Additionally, the *R*-(+) enantiomer of SKF-83959 in the PDB structure (PDB ID: 7jvp) share this similarity with the methyl group pointing towards this pocket (Fig. 6e). These A-868929 analogs also demonstrated high selectivity of D1R over D2R. Zhuang and coworkers indicated that D1 and D2Rs adopted distinct pockets in the extracellular region. By aligning of D1 and D2R structures, they found that the highly D1R selective SKF compounds extended closer to the ECL2 and clashed with I184^ECL2^ of D2R, while smaller S188^ECL2^ at the corresponding position of D1R pointed away from the ligand. Therefore, D1R has more space near the ECL2 region to accommodate large functional groups (Zhuang et al., 2021a). To evaluate if this rationale can be applied to explain the D1R selectivity of A-86929 analogs, we superimposed docking poses of these analogs in D1R with the D2R structure (PDB ID: 7jvr). As expected, the thiophene moiety and its substituents also clash with I184^ECL2^ of D2R (Fig. 7a-d). Additionally, I184^ECL2^ and F389^6.51^ form a ‘bottleneck’ which may further limit accessibility of the EBP of D2R for these A-86929 analogs (Fig. 7e)

**Figure 5.**
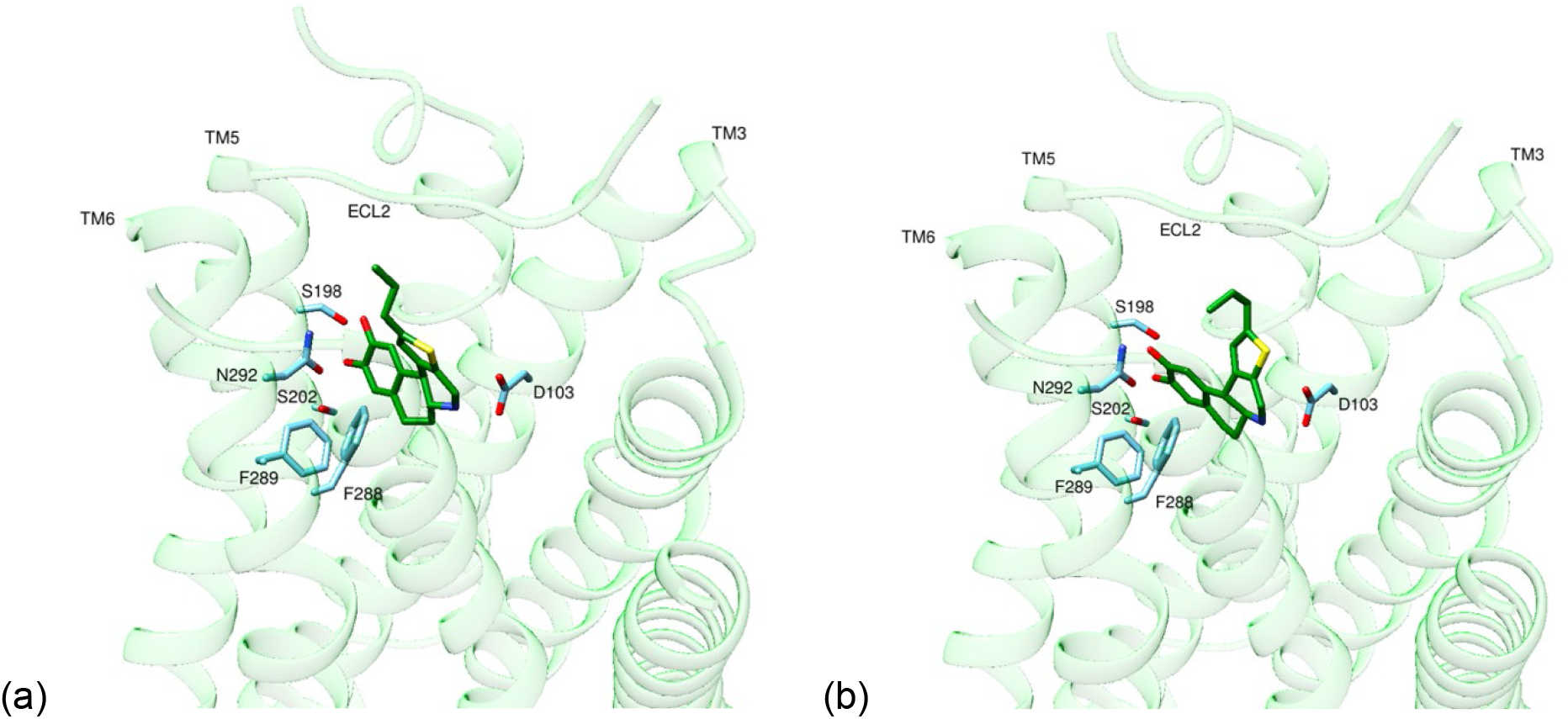
Putative binding poses of A-86929 and its enantiomer in D1R generated by molecular docking (D1R ribbon: green, D1R residues: blue, ligand: deep green). (a) A-86929 (b) (+)-enantiomer of A-86929

**Figure 6.**
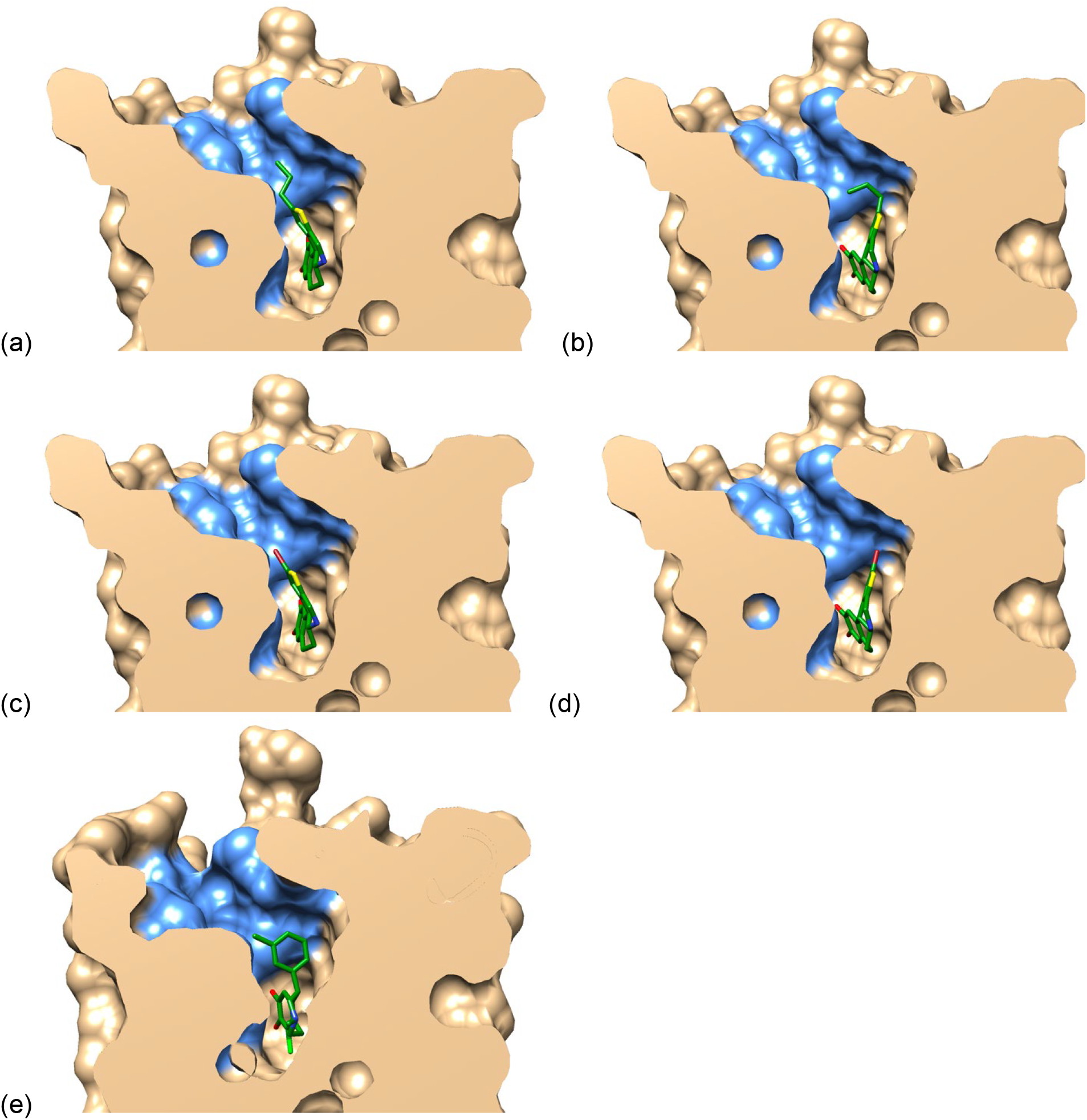
Clipping view of docking poses of A-86929 analogs and SKF-83959 PDB structure from TM1 of D1R (general D1R surface: tan, EBP surface: blue, ligands: deep green) (a) A-86929 (b) (+)-enantiomer of A-86929 (c) compound **1c** (d) compound **1d** (e) *R*-(+) SKF-83959

**Figure 7.**
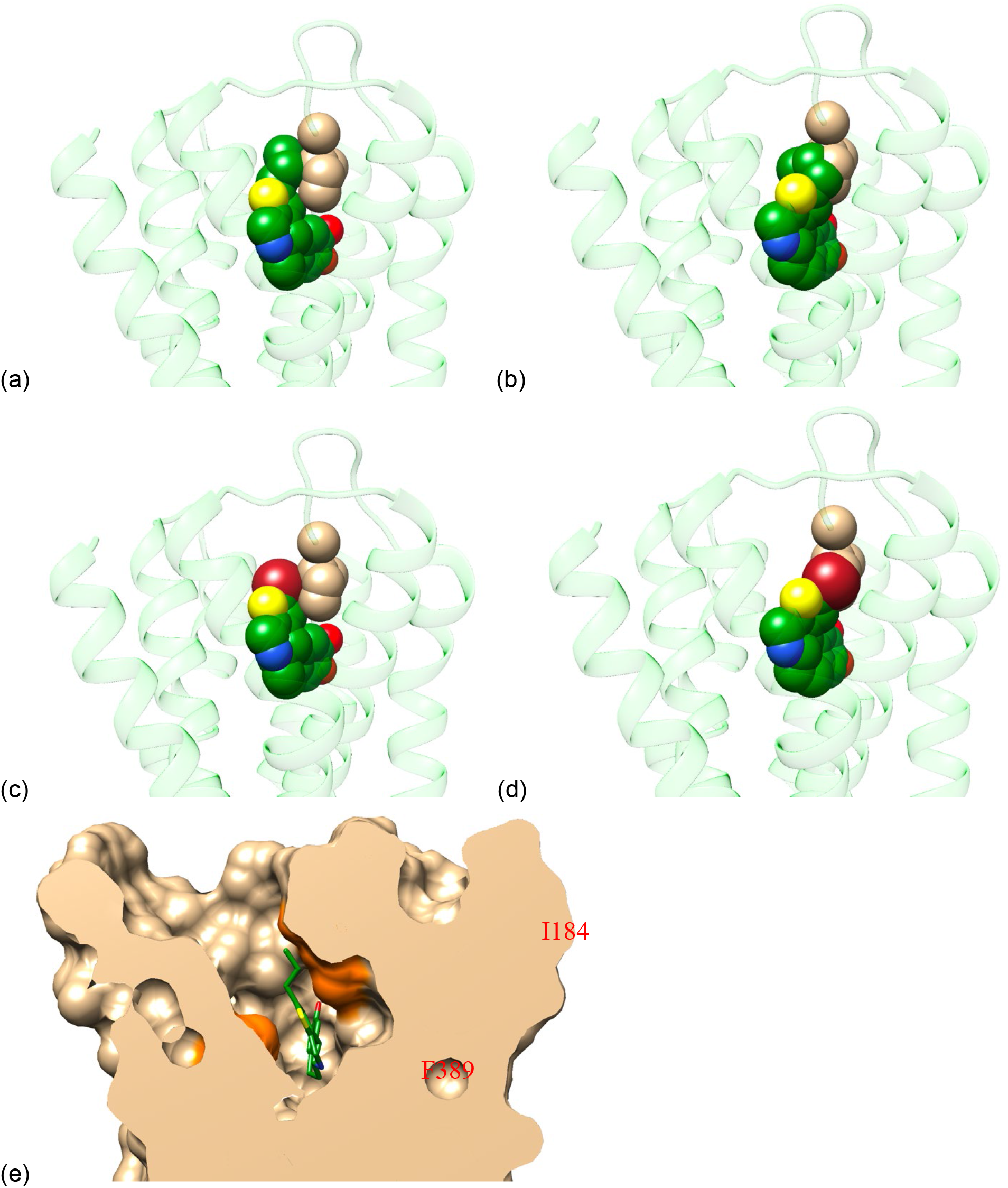
Superimposed D1R docking poses of A-86929 analogs with D2R structure (D1R is hidden in all figures). **a-d** Green ribbon: D2R, tan spheres: I184^ECL2^ of D2R, green spheres: carbon atoms of ligands, red spheres: oxygen atoms of ligands, blue spheres: nitrogen atoms of ligands, yellow spheres: sulfur atoms of ligands, brown spheres: bromine atoms of ligands. **e** clipping view of A-86929 analogs superimposed in D2R (general D2R surface: tan, I184 and F389 surfaces: orange, ligand: deep green) (a) A-86929 (b) (+)-enantiomer of A-86929 (c) compound **1c** (d) compound **1d** (e) A-86929

The amine group in dopamine has been shown to interact with the residue D103^3.32^ which is conserved in most Class A GPCRs, and that this residue is critical for receptor activation and signaling. Modifications at the amine group of many dopamine agonists either completely abolish or enhance signaling. For mixed dopamine agonists like apomorphine, modifications to the amine group (e.g., *N*-propylnorapomorphine (NPA)) renders it D2 selective owing to its picomolar potency at D2Rs (Park et al., 2020). With a methyl substitution on nitrogen of the azepane ring, SKF-82957 and SKF-83959 exhibit partial G protein biased signaling profile on D1R, while unsubstituted SKF-81297 is a signaling balanced D1R agonist (Conroy et al., 2015). Zhuang et al revealed the methyl group on nitrogen of SKF-83959 extended into a hydrophobic pocket surrounded by F288^6.51^, V317^7.39^, and W321^7.43^. Mutagenesis studies further suggested that F288^6.51^ and V317^7.39^ are crucial for β-arrestin activity of SKF compounds (Zhuang et al., 2021a). Consistently, when we added a methyl group to compound **1a**, we obtained a D1R G protein biased partial agonist **1j**. Our docking results suggested that this N-methyl group was accommodated within the hydrophobic pocket (Supplemental Fig. S1). Additionally, substitutions of the amino group for hydroxyl groups on dopamine-like fragments with ethyl (**3a**) or propyl (**3b**) linkers, which potentially alters H-bonding with D103^3.32^, gave some degree of arrestin bias at D2R vs D1R respectively, but reduced efficacy.

Given the critical role of ECL2 and amine group interactions, we hypothesized that smaller chemical scaffolds would allow for D1/D2R mixed agonism and reduce G protein activity, and give us an insight into the role of chemical interactions with ECL2 and the amine group potentially important for G protein vs arrestin signaling. All 2-aryl derivatives of PF-1437showed some degree of arrestin bias at D1Rs, except **2c** which showed G protein bias. On the other hand, **2a** and **c** displayed arrestin bias at D2Rs. To provide some structural insights into their activities, we also docked some of these molecules into D1R. Since these fragments are smaller in size, they may bind the receptor in multiple ways as conformational ensembles. We sampled 100 docking poses for each fragment and performed clustering analysis. The best pose of each cluster was selected to represent the cluster and two representative poses for each ligand are shown in Supplemental Fig. S2 a-g. The best representative poses for these active fragments always have interactions between the catechol moiety and N292^6.55^ or S198^5.42^. The other functionalized aromatic rings are placed close to hydrophobic residues identified above for compound **1j** (Fig. S1). The 2-aryl and 3-aryl fragments were largely D1 selective, possibly related to the D1R having more space near ECL2 allowing for larger functional groups. A combinatorial approach using 2-aryl fragments (**2a-h**) and the hydroxyl substituted dopamine analogs (**3a**, **b**) would be an interesting next step towards achieving more D1 selectivity and arrestin bias.

In conclusion, our SFSR study provides valuable insights into the chemical interactions that might drive G protein vs arrestin bias at D1Rs. Further SFSR on the small aryl scaffolds presented in this study, could potentially lead to identification of more efficacious and potent D1 selective arrestin biased agonists.

## ACKNOWLEDGEMENTS

We thank the financial support to this project from the Michael J. Fox Foundation (MJFF #11764 to N.U.) and the University of Florida Parkinson’s Moonshot Award (to N.U).

## METHODS

### cAMP GloSensor assay

To measure dopamine receptor mediated regulation of cAMP levels, HEK293T cells were co-transfected in a 1:1 ratio with either human D1 or D2Long receptor and a split-luciferase based cAMP biosensor (GloSensor, Promega, Durham NC). The next day, transfected cells were transferred to clear MEM media with 2% Fetal Bovine Serum (FBS) and 1X Glutamax, and plated in poly-D-lysine (Sigma Aldrich) coated 96-well white clear-bottom cell culture plates, at a density of 50,000 cells per 100 μl per well and incubated overnight. Next day, in a separate drug plate, serial drug dilutions ranging from 10^-3^M (1mM) to 10^-12^M (1pM) were prepared in fresh assay buffer (1X HBSS, 0.03% ascorbic acid, pH 7.4) such that the final concentrations would range from 10^-4^ to 10^-13^. Before adding drugs, 25 μl/well GloSensor reagent (4mM D-Luciferin, Cayman Chemicals) in assay buffer was added to each well. Plates were allowed to incubate for 2hrs in the dark at room temperature, and immediately afterwards, 10-20 μl assay buffer (to bring final volume to 50μl) and 5 μl of drugs or dopamine with concentrations corresponding to dose-response curves were added and allowed to incubate for an additional 5 minutes. To stimulate endogenous cAMP production (for D2R mediated inhibition) 5 μl of Forskolin (Sigma Aldrich) (1μM final concentration) was added after addition of drugs and incubated for an additional 5min. Luminescence intensity was quantified 15 minutes later using a Cytation 5 (BioTek) plate reader. The percent response was plotted as a function of drug concentration using Graphpad Prism 9 (Graphpad Software Inc., San Diego, CA).

### β-Arrestin Bioluminiscence resonance energy transfer (BRET) assay

To measure D1R- or D2R-mediated βarr2 recruitment, HEK293T cells were co-transfected in a 1:20 ratio with D1 or D2_Long_ receptor fused to C-terminal *renilla* luciferase (*R*Luc8 or 2), and a N-terminal Venus-tagged β-arrestin2. The next day, transfected cells were plated in poly-D-lysine coated 96-well white clear-bottom cell culture plates with clear MEM media + 2% FBS and 1X Glutamax at a density of 100,000 cells in 100 μl per well, and incubated overnight. Buffers used for the BRET assay and to dilute drugs were exactly the same as for the cAMP inhibition assay. Next day, media was decanted and cells were washed twice with assay buffer and 80 μl of assay buffer was added/well. The *R*Luc substrate, coelenterazine h (Cayman Chemicals, 5 μM final concentration), was added per well, and exactly 5 minutes later drugs were added at concentrations corresponding to the dose-response curves and allowed to incubate for 5 minutes. Luminescence at 485 nm and fluorescent eYFP emission at 530 nm were measured for 1 second per well using a Cytation 5 plate Reader (BioTek). The ratio of eYFP/*R*Luc was calculated per well and data are presented as percent of dopamine response. The percent response was plotted as a function of drug concentration using Graphpad Prism 9 (Graphpad Software Inc., San Diego, CA).

### Bias factor calculations

Using Graphpad Prism 9.0, transduction coefficients i.e log (τ/K_A_) were calculated for each drug at both pathways (Gαs and β-arrestin2) based on the Black and Leff operational model, where K_A_ is the equilibrium dissociation constant and τ is the agonist efficacy. Bias factors were calculated based on the method of Kenakin (Kenakin et al., 2012), where Δlog (τ/K_A_) for each drug was calculated in relation to the reference agonist dopamine, and the ΔΔlog (τ/K_A_) was calculated by subtracting the relative transduction coefficients [Δlog (τ/K_A_)] of the two pathways for each drug. A negative value represents arrestin bias whereas a positive value represents G protein bias.

### Molecular Docking

Molecular docking was implemented with Schrödinger Maestro. The D1R receptor structure PDB ID: 7jvq was chosen for this study since the bound cognate ligand, apomorphine, has a similar tetracyclic scaffold as the A-86929 analogs. To further evaluate if this structure was suitable for our ligands, enrichment analysis was performed. Ten tetracyclic molecules, which were experimentally tested as agonists for D1R, were selected and 50 decoys for each agonist were generated from the DUD-E database (Huang et al., 2006; Mysinger et al., 2012). Since the PDB structure corresponds to the G protein bound state, five antagonists were selected to test if docking can distinguish these antagonists from agonists. The total 515 ligands were docked into 7jvq structure and receiver operator characteristic area under the curve (ROCAUC) (Truchon and Bayly, 2007) value was calculated. The ROCAUC value was 0.99 and all true positives were recovered from the top 10% of the results. Additionally, all the agonists are ranked higher than four of the antagonists. These results indicated that this structure may be a suitable one to dock A-86929 analogs.

Glide XP mode was used to dock A-86929 analogs. The top scored poses were inspected and one pose was selected for each analog, in which the ligand had interactions with key residues identified by experimental mutational data of similar scaffold dihydrexidine on D1R (Mente et al., 2015). For the small fragments, we sampled 100 poses for each molecule with Glide SP mode. Then, the docking results were clustered (Ruvinsky and Kozintsev, 2005) and one pose represents each cluster. The top scored two representatives of each fragment were selected.

## Supplemental Figures and Tables

**Table S1.**
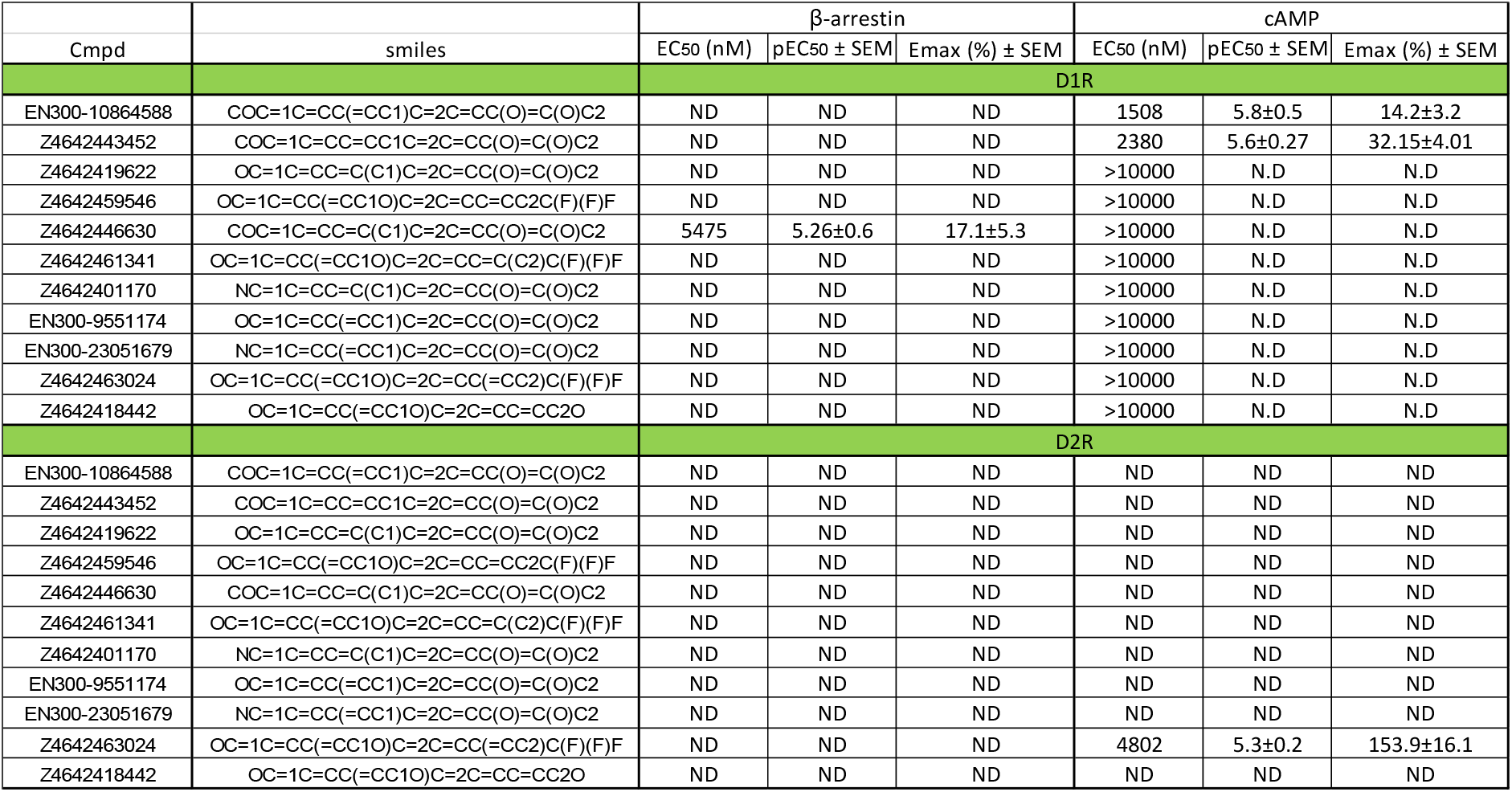
Agonist activities of 3-aryl catechol fragments EC_50_ and E_max_ values are from three independent experiments performed in duplicate or triplicate. E_max_ values are calculated as % response normalized to dopamine. ND - not detectable, N.D. not determined.

**Figure S1.**
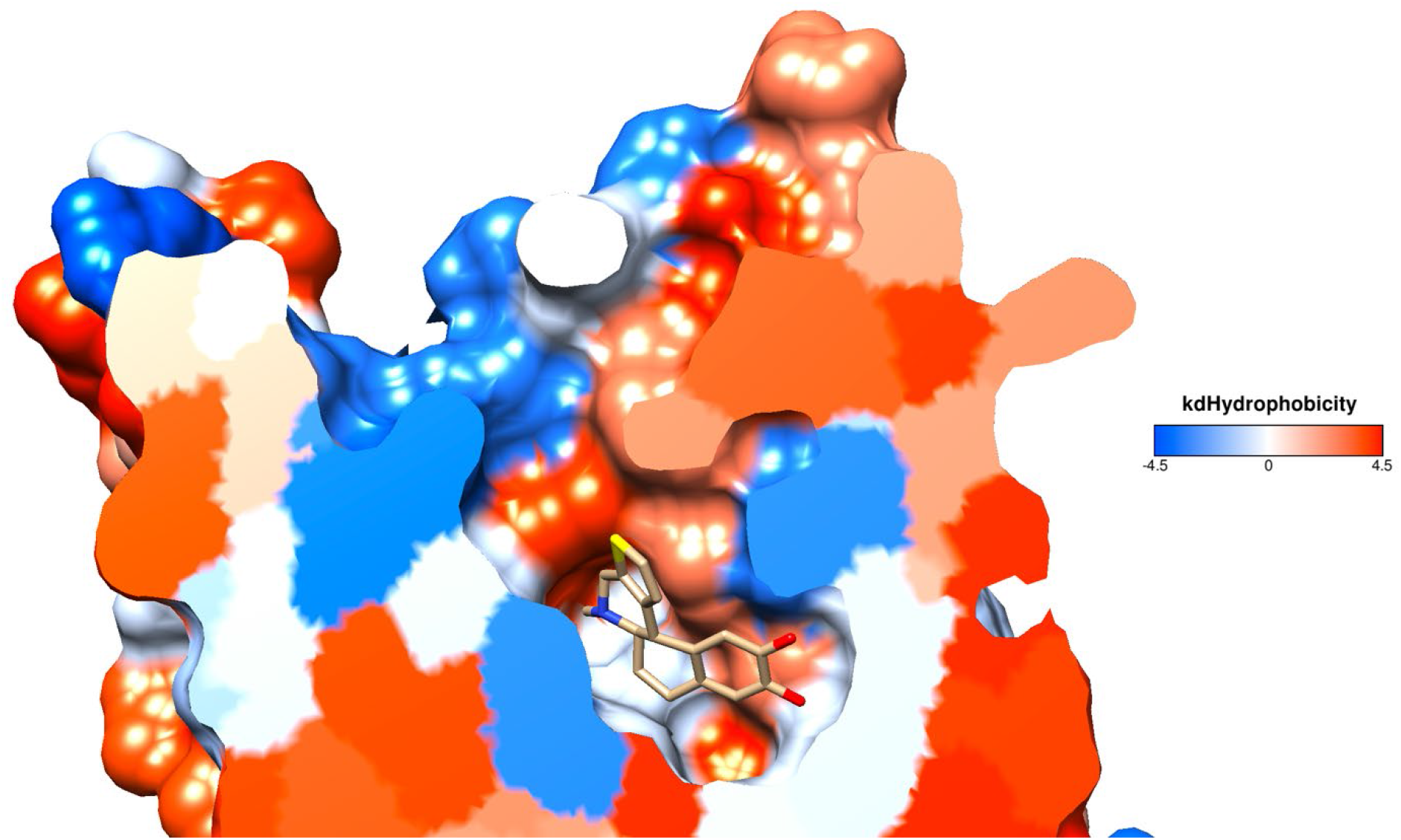
Clipping view of *5aR,* 11b*S* enantiomer of compound **1j** docking pose in D1R from TM3 (D1R surface color based on hydrophobicity scale of Kyte and Doolittle (Kyte and Doolittle, 1982), ligand: tan)

**Figure S2.**
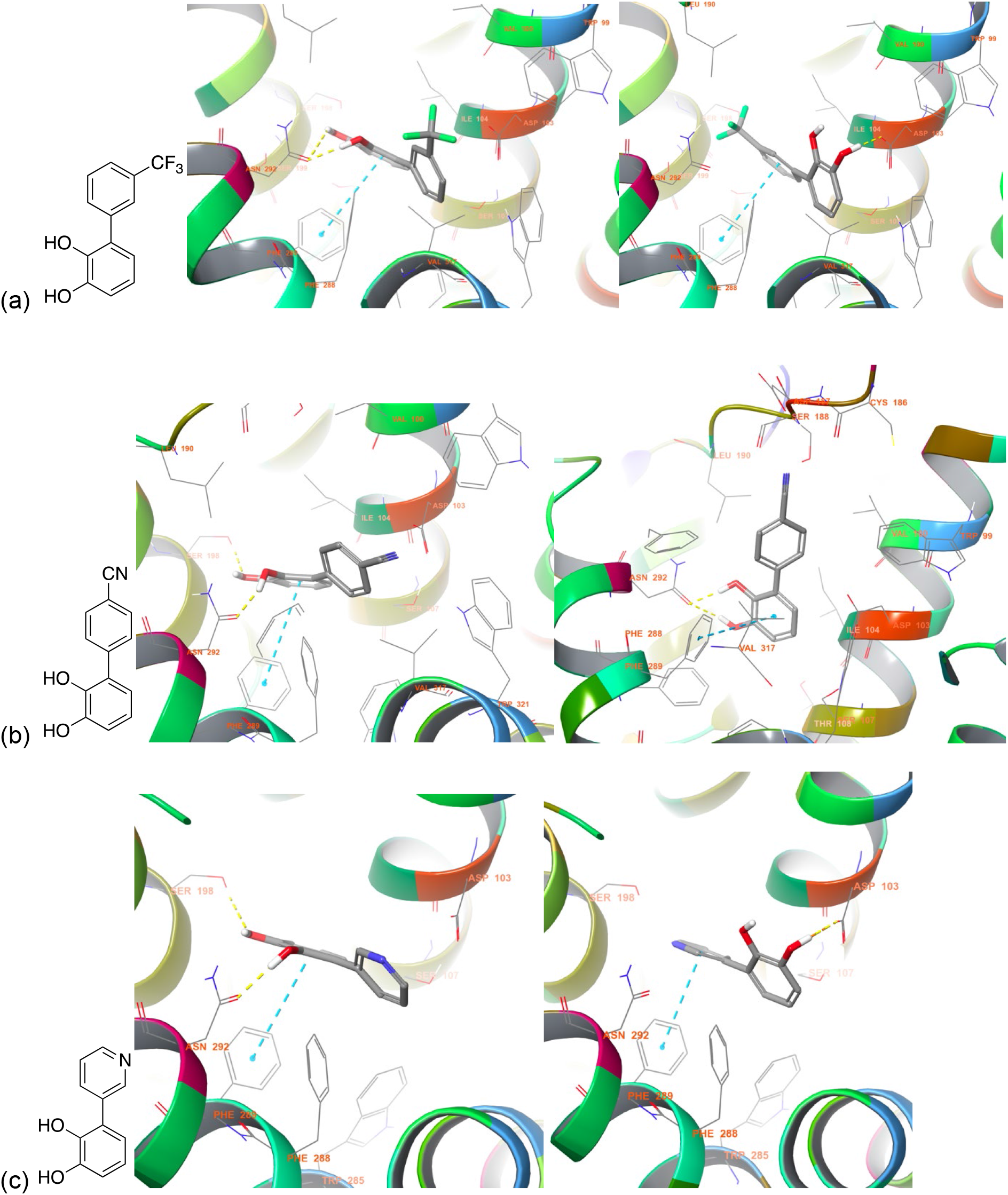
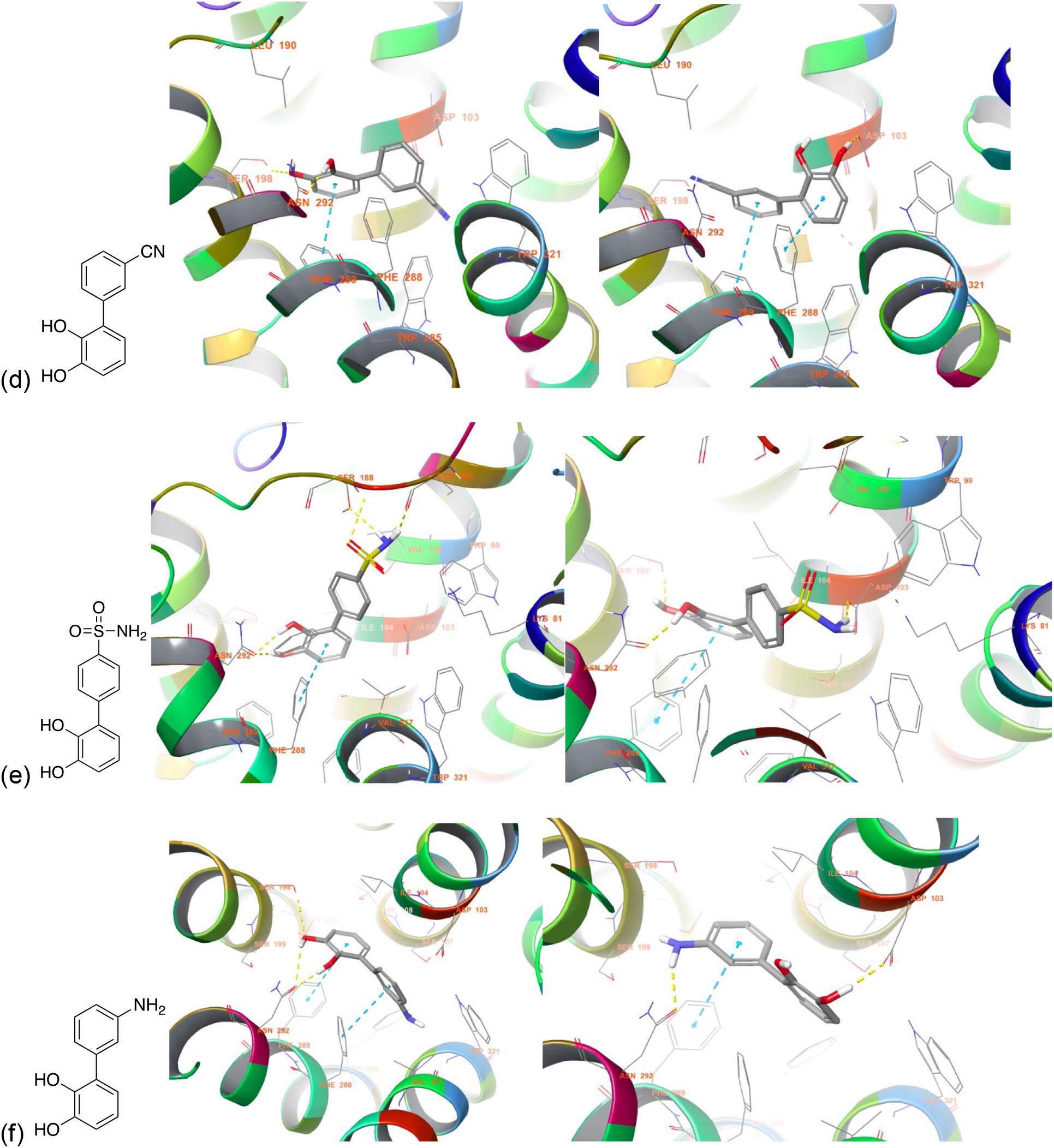
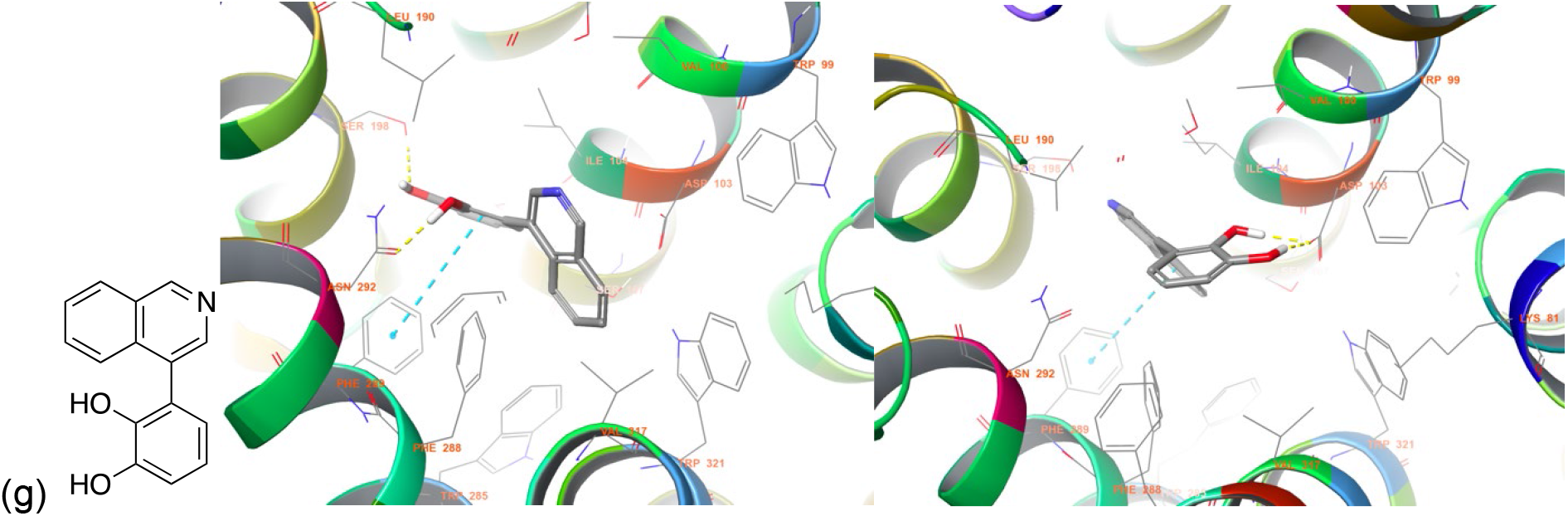
Top two representative docking poses for clusters of each fragment. (a) compound **2b** (b) compound **2c** (c) compound **2d** (d) compound **2e** (e) compound **2f** (f) compound **2g** (g) compound **2h**

## Notes

### Competing Interest Statement

The authors have declared no competing interest.

### Summary of Updates

1. errors in figures 3 and 4 have now been corrected 2. updated results and discussion 3. author affiliations updated

## REFERENCES

Allen, J.A., Yost, J.M., Setola, V., Chen, X., Sassano, M.F., Chen, M., Peterson, S., Yadav, P.N., Huang, X.P., Feng, B., et al. (2011). Discovery of beta-arrestin-biased dopamine D2 ligands for probing signal transduction pathways essential for antipsychotic efficacy. Proceedings of the National Academy of Sciences of the United States of America 108, 18488–18493.

Ballesteros, J.A., and Weinstein, H. (1995). [19] Integrated methods for the construction of three-dimensional models and computational probing of structure-function relations in G protein-coupled receptors.

Conroy, J.L., Free, R.B., and Sibley, D.R. (2015). Identification of G protein-biased agonists that fail to recruit beta-arrestin or promote internalization of the D1 dopamine receptor. ACS Chem Neurosci 6, 681–692.

Felsing, D.E., Jain, M.K., and Allen, J.A. (2019). Advances in Dopamine D1 Receptor Ligands for Neurotherapeutics. Current Topics in Medicinal Chemistry 19, 1365–1380.

Gray, D.L., Allen, J.A., Mente, S., O’Connor, R.E., DeMarco, G.J., Efremov, I., Tierney, P., Volfson, D., Davoren, J., Guilmette, E., et al. (2018). Impaired beta-arrestin recruitment and reduced desensitization by non-catechol agonists of the D1 dopamine receptor. Nat Commun 9, 674.

Hajra, S., and Bar, S. (2011). Catalytic enantioselective synthesis of A-86929, a dopamine D1 agonist. Chemical Communications 47, 3981–3982.

Huang, N., Shoichet, B.K., and Irwin, J.J. (2006). Benchmarking sets for molecular docking. Journal of Medicinal Chemistry 49, 6789–6801.

Jellinger, K.A., and Korczyn, A.D. (2018). Are dementia with Lewy bodies and Parkinson’s disease dementia the same disease? BMC Med 16, 34.

Kenakin, T., Watson, C., Muniz-Medina, V., Christopoulos, A., and Novick, S. (2012). A Simple Method for Quantifying Functional Selectivity and Agonist Bias. Acs Chemical Neuroscience 3, 193–203.

Kyte, J., and Doolittle, R.F. (1982). A SIMPLE METHOD FOR DISPLAYING THE HYDROPATHIC CHARACTER OF A PROTEIN. Journal of Molecular Biology 157, 105–132.

Martini, M.L., Liu, J., Ray, C., Yu, X., Huang, X.P., Urs, A., Urs, N., McCorvy, J.D., Caron, M.G., Roth, B.L., et al. (2019a). Defining Structure-Functional Selectivity Relationships (SFSR) for a Class of Non-Catechol Dopamine D1 Receptor Agonists. J Med Chem 62, 3753–3772.

Martini, M.L., Ray, C., Yu, X., Liu, J., Pogorelov, V.M., Wetsel, W.C., Huang, X.P., McCorvy, J.D., Caron, M.G., and Jin, J. (2019b). Designing Functionally Selective Noncatechol Dopamine D1 Receptor Agonists with Potent In Vivo Antiparkinsonian Activity. ACS Chem Neurosci 10, 4160–4182.

Mente, S., Guilmette, E., Salafia, M., and Gray, D. (2015). Dopamine D1 receptor-agonist interactions: A mutagenesis and homology modeling study. Bioorganic & Medicinal Chemistry Letters 25, 2106–2111.

Michaelides, M.R., Hong, Y., DiDomenico, S., Jr., Asin, K.E., Britton, D.R., Lin, C.W., Williams, M., and Shiosaki, K. (1995). (5aR,11bS)-4,5,5a,6,7,11b-hexahydro-2-propyl-3-thia-5-azacyclopent-1-ena[c]-phenanthrene-9,10-diol (A-86929): a potent and selective dopamine D1 agonist that maintains behavioral efficacy following repeated administration and characterization of its diacetyl prodrug (ABT-431). J Med Chem 38, 3445–3447.

Michaelides, M.R., Hong, Y., DiDomenico, S., Jr., Bayburt, E.K., Asin, K.E., Britton, D.R., Lin, C.W., and Shiosaki, K. (1997). Substituted hexahydrobenzo[f]thieno[c]quinolines as dopamine D1-selective agonists: synthesis and biological evaluation in vitro and in vivo. J Med Chem 40, 1585–1599.

Mysinger, M.M., Carchia, M., Irwin, J.J., and Shoichet, B.K. (2012). Directory of Useful Decoys, Enhanced (DUD-E): Better Ligands and Decoys for Better Benchmarking. Journal of Medicinal Chemistry 55, 6582–6594.

Park, H., Urs, A.N., Zimmerman, J., Liu, C., Wang, Q., and Urs, N.M. (2020). Structure-Functional-Selectivity Relationship Studies of Novel Apomorphine Analogs to Develop D1R/D2R Biased Ligands. ACS Med Chem Lett 11, 385–392.

Park, S.M., Chen, M., Schmerberg, C.M., Dulman, R.S., Rodriguiz, R.M., Caron, M.G., Jin, J., and Wetsel, W.C. (2015). Effects of beta-Arrestin-Biased Dopamine D2 Receptor Ligands on Schizophrenia-Like Behavior in Hypoglutamatergic Mice. Neuropsychopharmacology.

Rascol, O., Blin, O., Thalamas, C., Descombes, S., Soubrouillard, C., Azulay, P., Fabre, N., Viallet, F., Lafnitzegger, K., Wright, S., et al. (1999). ABT-431, a D1 receptor agonist prodrug, has efficacy in Parkinson’s disease. Annals of neurology 45, 736–741.

Rascol, O., Nutt, J.G., Blin, O., Goetz, C.G., Trugman, J.M., Soubrouillard, C., Carter, J.H., Currie, L.J., Fabre, N., Thalamas, C., et al. (2001). Induction by dopamine D1 receptor agonist ABT-431 of dyskinesia similar to levodopa in patients with Parkinson disease. Arch Neurol 58, 249–254.

Ruvinsky, A.M., and Kozintsev, A.V. (2005). New and fast statistical-thermodynamic method for computation of protein-ligand binding entropy substantially improves docking accuracy. Journal of Computational Chemistry 26, 1089–1095.

Shepherd, G.M. (2013). Corticostriatal connectivity and its role in disease. Nat Rev Neurosci 14, 278–291.

Shiflett, M.W., and Balleine, B.W. (2011). Molecular substrates of action control in cortico-striatal circuits. Prog Neurobiol 95, 1–13.

Smith, K.S., and Graybiel, A.M. (2016). Habit formation. Dialogues Clin Neurosci 18, 33–43.

Sodergren, M.J., Alonso, D.A., and Andersson, P.G. (1997). Readily available nitrene precursors increase the scope of Evans’ asymmetric aziridination of olefins. Tetrahedron-Asymmetry 8, 3563–3565.

Surmeier, D.J., Obeso, J.A., and Halliday, G.M. (2017). Selective neuronal vulnerability in Parkinson disease. Nat Rev Neurosci 18, 101–113.

Truchon, J.F., and Bayly, C.I. (2007). Evaluating virtual screening methods: Good and bad metrics for the “early recognition” problem. Journal of Chemical Information and Modeling 47, 488–508.

Urban, J.D., Clarke, W.P., von Zastrow, M., Nichols, D.E., Kobilka, B., Weinstein, H., Javitch, J.A., Roth, B.L., Christopoulos, A., Sexton, P.M., et al. (2007). Functional selectivity and classical concepts of quantitative pharmacology. J Pharmacol Exp Ther 320, 1–13.

Urs, N.M., Bido, S., Peterson, S.M., Daigle, T.L., Bass, C.E., Gainetdinov, R.R., Bezard, E., and Caron, M.G. (2015). Targeting beta-arrestin2 in the treatment of L-DOPA-induced dyskinesia in Parkinson’s disease. Proceedings of the National Academy of Sciences of the United States of America 112, E2517–2526.

Urs, N.M., Gee, S.M., Pack, T.F., McCorvy, J.D., Evron, T., Snyder, J.C., Yang, X., Rodriguiz, R.M., Borrelli, E., Wetsel, W.C., et al. (2016). Distinct cortical and striatal actions of a beta-arrestin-biased dopamine D2 receptor ligand reveal unique antipsychotic-like properties. Proceedings of the National Academy of Sciences of the United States of America 113, E8178–E8186.

Wang, P.Y., Felsing, D.E., Chen, H.Y., Raval, S.R., Allen, J.A., and Zhou, J. (2019). Synthesis and Pharmacological Evaluation of Noncatechol G Protein Biased and Unbiased Dopamine D1 Receptor Agonists. Acs Medicinal Chemistry Letters 10, 792–799.

Wickens, J.R., Horvitz, J.C., Costa, R.M., and Killcross, S. (2007). Dopaminergic mechanisms in actions and habits. J Neurosci 27, 8181–8183.

Wood, J., and Ahmari, S.E. (2015). A Framework for Understanding the Emerging Role of Corticolimbic-Ventral Striatal Networks in OCD-Associated Repetitive Behaviors. Front Syst Neurosci 9, 171.

Xiao, P., Yan, W., Gou, L., Zhong, Y.N., Kong, L., Wu, C., Wen, X., Yuan, Y., Cao, S., Qu, C., et al. (2021). Ligand recognition and allosteric regulation of DRD1-Gs signaling complexes. Cell.

Yang, Y., Lee, S.M., Imamura, F., Gowda, K., Amin, S., and Mailman, R.B. (2018). D1 dopamine receptors intrinsic activity and functional selectivity affect working memory in prefrontal cortex. Mol Psychiatry.

Yin, H.H., Knowlton, B.J., and Balleine, B.W. (2006). Inactivation of dorsolateral striatum enhances sensitivity to changes in the action-outcome contingency in instrumental conditioning. Behav Brain Res 166, 189–196.

Zhang, J., Xiong, B., Zhen, X.C., and Zhang, A. (2009). Dopamine D-1 Receptor Ligands: Where Are We Now and Where Are We Going. Medicinal Research Reviews 29, 272–294.

Zhuang, Y., Xu, P., Mao, C., Wang, L., Krumm, B., Zhou, X.E., Huang, S., Liu, H., Cheng, X., Huang, X.P., et al. (2021a). Structural insights into the human D1 and D2 dopamine receptor signaling complexes. Cell.

Zhuang, Y.W., Krumm, B., Zhang, H.B., Zhou, X.E., Wang, Y., Huang, X.P., Liu, Y.F., Cheng, X., Jiang, Y., Jiang, H.L., et al. (2021b). Mechanism of dopamine binding and allosteric modulation of the human D1 dopamine receptor. Cell Research 31, 593–596.

